# Stronger together: isograft fusion accelerates recovery and may facilitate heat-evolved symbiont transmission in bleached adult corals

**DOI:** 10.64898/2026.08.11.744325

**Authors:** Bede G. Johnston, Catalina Parra V, Matthew R. Nitschke, Wing Yan Chan, Madeleine J.H. van Oppen

**Author notes:** Corresponding author: Mailing address: PMB 3, Townsville MC, 4810 QLD, Australian Institute of Marine Science, 1526 Cape Cleveland Road, Cape Cleveland, Townsville, QLD 4810, Australia.

## Abstract

Experimentally evolved, heat-tolerant algal symbionts (heat-evolved; HE) offer a promising means of enhancing coral holobiont thermotolerance under rapidly warming oceans. However, translating their benefits into restoration practices requires scalable delivery methods. Coral tissue fusion may provide one such pathway by facilitating HE symbiont transfer to wild corals; however, its feasibility remains largely untested. As an initial test, we paired adult isografts of *Galaxea fascicularis* and *Psammocora columna* hosting HE *Cladocopium proliferum* (SS8) with chemically bleached, SS8-naive recipients. Fusion was first observed after three days in *G. fascicularis* and nine days in *P. columna*. In both species, fusion was followed by increased pigmentation and photochemical efficiency at the recipient’s fusion interface relative to distal tissue and unfused controls. After ∼50 days, SS8 was detected at low levels (<3.5%) in 15/19 fused *G. fascicularis* recipients, although detection was also common among unfused horizontal-transmission controls maintained in the same water column (13/18). These findings provide the first empirical evidence that conspecific coral tissue fusion is associated with localised physiological recovery and can coincide with HE symbiont acquisition, while highlighting the need to distinguish tissue-mediated transfer from background horizontal transmission. Fusion may therefore represent a complementary pathway for beneficial symbiont delivery in assisted-evolution frameworks.

## Introduction

The success of coral reef ecosystems is underpinned by the symbiotic relationship between reef-building corals and endosymbiotic dinoflagellates (family Symbiodiniaceae) [1]. Residing within coral gastrodermal cells, these photosynthetic, microalgal symbionts provide the coral host with most of its nutritional requirements by converting inorganic carbon and host waste products into energy-rich photosynthates [2–6]. Although highly efficient, this metabolic exchange can become unstable under stressful environmental conditions, such as elevated temperatures or excessive light exposure, leading to host energetic stress, and, if stress persists, degradation or expulsion of the algal symbionts — i.e., coral bleaching [7, 8].

The ability of corals to tolerate and recover from environmental stress is strongly influenced by the physiological performance of their resident Symbiodiniaceae [7, 9, 10]. For example, corals associated with naturally thermally tolerant symbionts, including some *Durusdinium* spp., often exhibit improved bleaching tolerance under elevated temperatures [10–15]. However, associations with naturally thermotolerant symbionts have been linked to context-dependent physiological trade-offs ([16–19], but see [10, 20, 21]) and may still fail to prevent bleaching during severe marine heatwaves [22]. Such limitations have motivated the development of symbiont manipulation approaches that enhance coral thermal tolerance while minimising trait trade-offs.

One promising approach is the experimental evolution of Symbiodiniaceae under thermal selection, in which cultured symbiont populations are subjected to prolonged elevated temperatures to generate strains with improved heat tolerance [23, 24]. Using a thermal ratchet design [25], Chakravarti *et al.* [26], [27] generated several heat-evolved (HE) strains of *Cladocopium proliferum* that exhibited enhanced growth and photochemical efficiency under acute heat stress relative to their wild-type (WT) counterparts. For some HE strains, these benefits have also been shown to extend to the coral holobiont, with no observable trait trade-offs [28–30].

The integration of HE symbionts into reef restoration practices requires scalable delivery methods that promote reliable uptake and persistence within coral hosts [24, 31]. Adult coral tissue fusion may offer one such pathway by creating direct tissue continuity between colonies through which symbionts could potentially move across the fusion boundary. However, whether fusion can facilitate symbiont movement between adult corals remains largely untested, particularly for experimentally evolved heat-tolerant strains.

Coral tissue fusion is a well-documented and ecologically important life-history strategy observed across multiple life stages. It occurs frequently among gregariously settled larvae [32–35], closely related juveniles [36–38], and genetically identical adult fragments [39–41], with some evidence also reported for allogeneic fusion between genetically distinct conspecifics [42, 43]. Fusion has been shown to confer a number of physiological benefits, including accelerated growth, expanded spatial coverage, and earlier onset of sexual maturity [36, 39, 40]. These benefits are thought to arise, at least in part, through integration of the colonies’ gastrovascular systems (GVS), which support gas exchange, extracellular digestion, waste removal, and the distribution of dissolved and particulate organic matter throughout the colony [44].

Beyond organic matter and nutrients, intact Symbiodiniaceae cells have been observed circulating within the coral coelenteron (gastrovascular cavity; GVC) [44, 45]. Although such cells are often presumed to be digested or expelled via the polyp mouth [3, 46, 47], gastrovascular integration following fusion could allow donor-derived symbionts to enter the GVC of an adjacent recipient colony. Once there, symbionts may be captured and internalised by the ciliated endodermal cells as observed in Hirose *et al.* [48]. If retained, these cells could contribute to symbiont establishment within recipient tissues, providing a biologically plausible pathway through which fusion could promote symbiont uptake in coral hosts.

This study investigates whether adult isograft tissue fusion between a donor fragment hosting HE *C. proliferum* (strain SS8) and a chemically bleached recipient fragment can facilitate the detectable acquisition of SS8. To achieve this, donor and recipient fragments were generated within each of two coral species, *Galaxea fascicularis* and *Psammocora columna*. Fragments were cut to expose adjacent tissue margins, positioned in direct contact with conspecific paired fragments, and maintained for ∼50 days, during which fusion progression, tissue repigmentation, and photochemical efficiency were regularly assessed. At the conclusion of the experiment, tissue samples were collected from defined positions relative to the fusion boundary and analysed using Internal Transcribed Spacer 2 (ITS2) amplicon sequencing to detect SS8 where strain-level resolution was possible. Together, this approach provides a controlled first test of whether coral tissue fusion can facilitate symbiont acquisition between adult coral fragments.

## Materials and Methods

### Collection and maintenance of Galaxea fascicularis and Psammocora columna colonies

Three colonies of *G. fascicularis* (herein referred to as gA, gB, or gC) were collected from Davies Reef in the central Great Barrier Reef (GBR) on October 29, 2021, at depths of 5-10 m (collection permit: G12/35236.1). In addition, four colonies of *P. columna* (pA, pB, pC, and pD) were collected from Fantome Reef in the central GBR on May 25, 2022, at a depth of ∼3 m (collection permit: G21/38062.1). Distances between individual colonies were > 5 m and are thus assumed to represent distinct genotypes. All colonies were transported to the National Sea Simulator at the Australian Institute of Marine Science, Townsville, and acclimated for two weeks. Following acclimation, *G. fascicularis* colonies were sectioned into 4–5 polyp fragments using a diamond blade band saw (Gryphon C-40 Bandsaw) and affixed to 22 × 30 mm PVC tiles using Gorilla Super Glue while *P. columna* colonies were cut into ∼3 cm² fragments and affixed to aragonite frag plugs (Large OW100LCFP, Aquasonic, Wauchope, Australia). Fragments were maintained in 2 µm filtered seawater (FSW) with a continuous FSW turnover rate of 5 L h⁻¹. Corals were maintained under an 11-hour photoperiod (06:50– 17:50) provided by Hydra 64HD LED lights (Aqua Illumination, Bethlehem, PA, USA), delivering within-tank light intensities ranging from 102–180 µmol photons m⁻² s⁻¹. Corals were fed daily with *Artemia* nauplii (0.5 nauplii mL⁻¹). Filamentous algae and biofilm were manually removed from tiles and plugs 1–2 times per week. Water temperature was maintained at 26.7–27.0°C. Water movement and aeration were provided using air stones.

### Heat-evolved symbiont (SS8)

The heat-evolved symbiont used was *Cladocopium proliferum* (SCF 055.01.08; referred to as SS8 as per [28–31]). Cultures of SS8 were cultivated at 31°C under a light intensity of 40-70 µmol photons m⁻² s⁻¹ with a 12:12 hour light-dark cycle. The cultivation medium used was 1% standard IMK culture medium (Nihon Pharmaceutical Co.). Cultures were refreshed every four weeks, involving the replacement of more than 50% of the existing culture medium with fresh IMK media.

### Generation of coral donors and recipients

SS8-*G. fascicularis* and SS8-*P. columna* donors were generated via chemical bleaching and subsequent inoculation with cultured SS8. The chemical bleaching protocol was adapted from Scharfenstein *et al.* [49] and refined through preliminary trials to balance sufficient symbiont reduction with host tissue integrity and survivorship. The selected menthol-diuron exposure regime induced visible paling and reduced photochemical signal while avoiding extensive tissue loss or mortality. Briefly, a subset of coral fragments from both species was incubated in 5 L of FSW supplemented with menthol (0.39 mM) and diuron (0.13 µM) for eight hours daily (08:30–16:30) during which gentle aeration was provided using air stones. Following the incubation period, menthol-diuron-spiked FSW was removed, and fragments were returned to FSW with regular turnover (5 L per hour) for 16 hr (16:30-08:30). This cycle was repeated for four consecutive days followed by a three-day recovery period, together, constituting one round of chemical bleaching. For *P. columna*, one round was applied, comprising four treatment days followed by three recovery days. For *G. fascicularis*, two complete rounds were applied consecutively, comprising eight treatment days and two three-day recovery periods, for a total bleaching schedule of 14 days. The number of rounds was determined from preliminary trials and visual assessment under a high-power stereomicroscope, with *P. columna* reaching sufficient bleaching after one round and *G. fascicularis* requiring a second round. No feeding occurred during the treatment or recovery periods.

Chemically bleached *G. fascicularis* and *P. columna* fragments were then inoculated daily with cultured SS8 (final cell density: 10,000 cells mL⁻¹) in line with previous coral– Symbiodiniaceae inoculation approaches [49]. Inoculations were performed during the culture’s peak motile period (∼1 hr after the culture incubator lights turned on) over a seven-day period. Corals were then allowed to repigment for seven weeks. Coral recovery was monitored via image-based red, green, blue (RGB) reflectance analysis and dark-adapted maximum quantum yield of photosystem II (*F_v_/F_m_*) (details provided below). ITS2 community composition analysis confirmed the presence of SS8 in donor fragments (details provided below). To generate coral recipients for subsequent tissue fusion, the remaining subset of *G. fascicularis* and *P. columna* fragments underwent the same chemical bleaching procedure, timed such that they would be sufficiently bleached by the end of the donors’ seven-week recovery period.

### Coral tissue fusion

#### Experimental treatments

To generate donor and recipient isografts, corals were first detached from their PVC tiles or aragonite plugs using a diamond-blade band saw (Gryphon C-40). Recipient fragments were cut perpendicularly to the base. Each fragment was first halved, and one half was then divided again to produce three pieces: one larger half-fragment and two smaller quarter-fragments (Fig. 1A). The larger piece was assigned to the fused recipient (FR) treatment. The two smaller pieces remained unfused and were allocated to control treatments: a horizontal transmission control (HT) placed in the same tank as SS8 donor fragments and fused donor–recipient pairs; and a symbiont-exclusion control (EC) maintained in a separate tank without exposure to external SS8 symbionts. Donor colonies were similarly bisected to produce two half-fragments: one assigned to the fused donor (FD) treatment and the other to the unfused donor (UD) control which was maintained in the same tank but kept physically separate from all recipient fragments.

**Fig 1.**
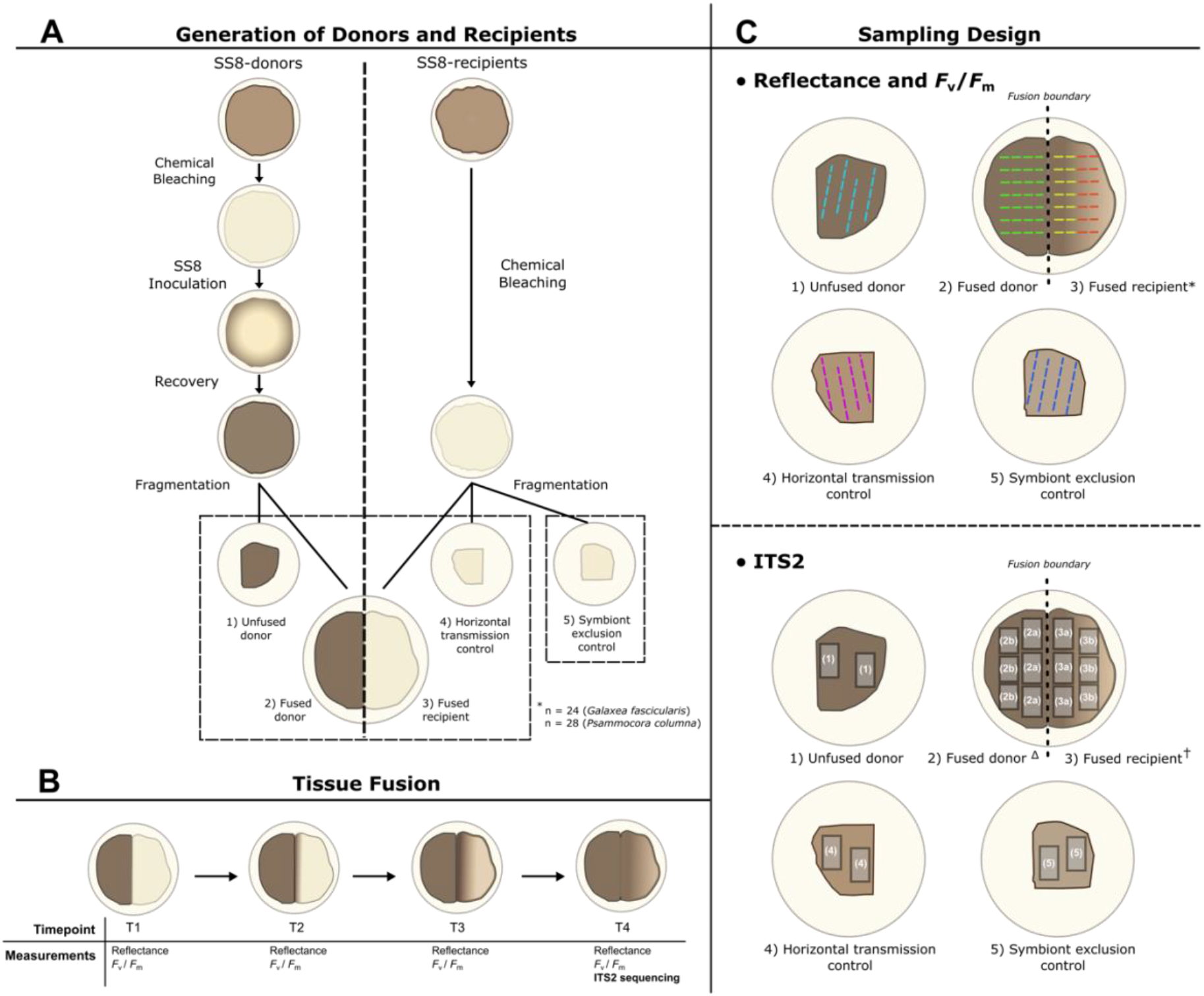
Schematic overview of the experimental design used to assess coral-to-coral transmission of the heat-evolved *Cladocopium proliferum* strain SS8 in *Psammocora columna* and *Galaxea fascicularis*. **(A)** SS8-donor corals were generated via chemical bleaching and inoculation with SS8, followed by a recovery period. SS8-recipient corals were generated by chemically bleaching fragments from the same source colonies immediately prior to the experiment. Donor and recipient fragments were cut using a band saw and allocated to one of five treatments: (1) unfused donor (UD), donor fragments maintained independently; (2) fused donor (FD), representing the donor side of fused donor-recipient pairs; (3) fused recipient (FR), representing the recipient side of fused donor-recipient pairs; (4) horizontal transmission control (HT), recipient fragments maintained in shared tanks alongside treatments 1–3; and (5) symbiont-exclusion control (EC), recipient fragments maintained in isolated tanks devoid of external SS8 sources. Tank allocation is indicated by hatched boxes. **(B)** Coral photophysiology, measured as maximum quantum yield of photosystem II (*F_v_*/*F_m_*), and reflectance were measured at multiple timepoints throughout the experiment to assess spatial and temporal patterns of recipient recovery. Endpoint tissue samples were collected after ∼50 days for Internal Transcribed Spacer 2 (ITS2) amplicon sequencing. **(C)** Sampling design for reflectance, *F_v_*/*F_m_*, and ITS2 analysis. *Top:* For UD, HT, and EC fragments, a minimum of four transects were drawn across the tissue surface in ImageJ [50]. For fused donor–recipient pairs, at least seven transects were positioned perpendicular to the fusion line, with the midpoint of each transect centred over the line. *For the fused recipient, transects were subdivided into two spatial zones: recipient-boundary (yellow) and recipient-far (red), allowing for the detection of localised variation in recovery. *Bottom:* Endpoint tissue sampling for ITS2 sequencing was similarly spatially structured. For UD, HT, and EC fragments, two technical replicates were collected from each fragment, one from the top surface and one from the side. For fused treatments 2^Δ^ (FD) and 3^†^ (FR), samples were collected from defined spatial zones: (a) tissue adjacent to the fusion line (termed donor boundary and recipient boundary), and (b) tissue located distally from the line (donor far and recipient far), to capture spatial variation in symbiont identity across the fusion interface. Technical replicates sharing the same number or number–letter identifier were extracted separately, and the resulting DNA extracts were pooled before PCR amplification and library preparation.

Immediately following sectioning, donor and recipient fragments were placed in their respective tanks and allowed to recover for four days before fusion setup. This recovery period was included to reduce the likelihood that symbionts released during cutting and acute tissue damage would contribute to waterborne acquisition during the initial fusion period. After this recovery period, fragments of *G. fascicularis* and *P. columna* from the same genotype— assigned to the FR and FD treatments—were carefully aligned on a new PVC tile or aragonite plug. To maximise tissue contact, the freshly cut lateral margins—comprising both tissue and underlying skeleton—were positioned flush against one another and secured using super glue to promote tissue fusion (see Supplementary Information Fig S1).

#### Tank allocation

A total of four 50 L tanks were used for the experiment, with two tanks allocated to each species. For each species, one tank housed all SS8-exposed treatments, including FR, FD, HT, and UD fragments, while the second tank housed EC fragments that were not exposed to external SS8 symbionts. Thus, the *G. fascicularis* and *P. columna* experiments each comprised one SS8-exposed tank and one EC tank.

Each donor–recipient pairing and its corresponding unfused control fragments were derived from the same source genotype and treated as a biological replicate. For *G. fascicularis*, each treatment included eight replicate fragment sets per genotype across three genotypes (*n* = 24 per treatment), while for *P. columna*, each treatment included seven replicate fragment sets per genotype across four genotypes (*n* = 28 per treatment). Corals were maintained in their respective tanks for ∼50 days, with physiological measurements taken at regular timepoints throughout the experiment and ITS2 samples collected at its conclusion (Table 1). Tank conditions and coral husbandry followed procedures as described above.

**Table 1:**
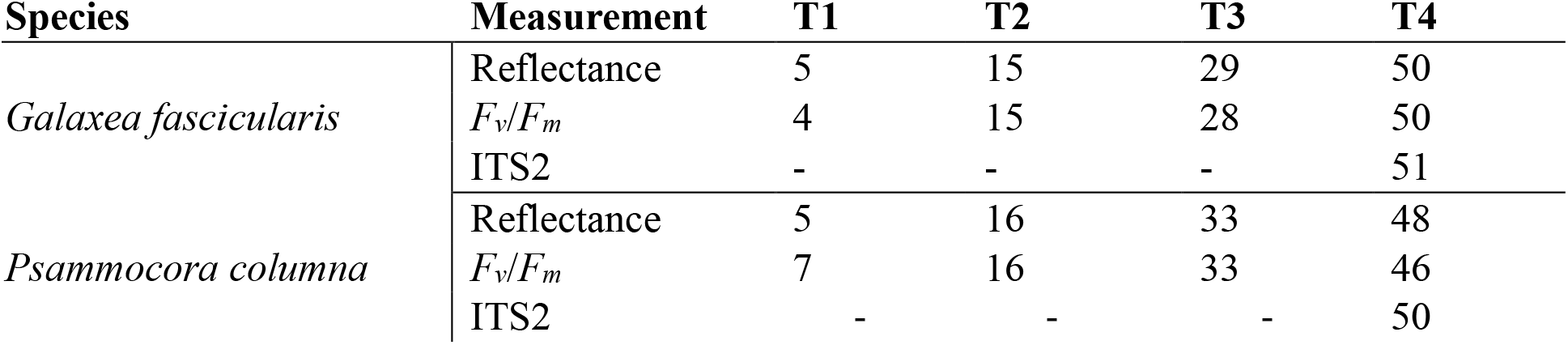
Sampling schedule for physiological and molecular measurements in *Galaxea fascicularis* and *Psammocora columna.* Timepoints (T1-4) are reported as days post-fusion setup (DPF) with 0 DPF defined as the day donor and recipient fragments were placed in direct contact to initiate fusion.

### Coral reflectance analysis

Coral red, green, and blue (RGB) reflectance values, used here as a proxy for bleaching severity, were quantified in ImageJ [50] using a transect-based approach. Photographs of each coral fragment were taken at designated timepoints (Table 1) using a Nikon D810 camera set to aperture priority mode (f/8, ISO 640, 1/80 s), with flash illumination provided by a Godox V860III Speedlite. Standardised lighting and positioning were maintained across timepoints to minimise variation. To calculate RGB reflectance for UD, HT, and EC treatments, a minimum of four transects were placed across each fragment (Fig. 1C; dashed lines). Each transect spanned ∼75% of the fragment’s width to capture intra-fragment variation while excluding areas affected by uneven lighting (due to fragment geometry). RGB values were extracted from every pixel along each transect and averaged to yield mean values per transect for each RGB channel. One pixel is equivalent to 0.035 mm.

For FR and FD, a minimum of seven transects were placed per fused pair, each spanning 60– 75% of the total fused fragment width and oriented perpendicular to the fusion line (Fig. 1C). Each transect was centred precisely over the visible fusion interface to enable targeted analysis of both donor and recipient tissue along a consistent spatial axis. To minimise potential misclassification at the immediate fusion line where tissue origin may be ambiguous, the central 0–5 pixels on either side of the midpoint were excluded from analysis (∼0.35 mm total). The remaining pixels were categorised into three spatial zones: (a) recipient boundary (5–55 pixels left of centre), (b) recipient far (55–105 pixels left of centre), and (c) donor (5–105 pixels right of centre). Mean reflectance values were calculated separately for each zone to capture localised variation across the fusion interface. RGB values were calibrated in ImageJ using six reference standards with known reflectance values. For each RGB channel, a 4th-degree polynomial regression was applied to generate calibration curves. To account for minor lighting variation across timepoints, separate calibration curves were generated for each sampling interval. All reflectance values reported in this study are based on these calibrated measurements.

### Maximum quantum yield of PSII (F_v_/F_m_)

The photochemical efficiency of coral fragments across all treatments was assessed using an imaging pulse-amplitude modulation fluorometer (IMAGING-PAM, Maxi version, Walz) (see Supplementary Information Fig S2 for representative image). Corals were dark-adapted for 15 minutes under low-light conditions (<5 μmol photons m⁻² s⁻¹) prior to measurement. Measurements of maximum quantum yield of photosystem II (*F_v_*/*F_m_* = [*F_m_−F_0_*]/*F_m_*) were conducted between 11:00 – 12:00 at designated timepoints (see Table 1). The same transect-based approach described for reflectance analysis was applied to extract *F_v_*/*F_m_* values across each coral fragment (Fig. 1C). Imaging-PAM parameters were set as follows: measuring light intensity = 4, gain = 3, frequency = 4, damping = 2. Data were collected via ImagingWin software (V2.32 FW Multi RGB; Walz GmbH, Effeltrich, Germany).

Physiological measurements were conducted at four approximately matched timepoints, with slight differences between species and measurement types (Table 1). For *G. fascicularis*, reflectance was measured at 5, 15, 29, and 50 DPF, *F_v_*/*F_m_* at 4, 15, 28, and 50 DPF, and ITS2 samples were collected at 51 DPF. For *P. columna*, reflectance was measured at 5, 16, 33, and 48 DPF, *F_v_*/*F_m_* at 7, 16, 33, and 46 DPF, and ITS2 samples were collected at 50 DPF.

### Macro-photography

A series of high-resolution close-up images (0.63× to 4× magnification) of all fusion pairings were captured using a Leica MZ10F stereo microscope equipped with a DFC450 C digital camera. These images were taken to provide a detailed time series documenting the progression of fusion events, including the timepoint when fusion was first confirmed. Fusion was deemed successful when the tissues of adjoining fragments had visibly merged at the interface and the origin of each tissue could no longer be distinguished. Illumination for imaging was provided by an EL6000 external light source and a dual-arm LED gooseneck.

### Sampling design

Per fragment pairing (inclusive of treatments 1-5), a total of seven sample types were collected: (1) UD, (2a) FD boundary, (2b) FD far, (3a) FR boundary, (3b) FR far, (4) HT, and (5) EC (see Fig. 1C). To generate sample types 2a/b and 3a/b, the fused coral pair was first sectioned along the visible fusion line to separate the donor and recipient fragments (see Supplementary Information Fig S3). To minimise cross-contamination between donor and recipient tissues, ∼1 mm of tissue and underlying skeleton was shaved from the adjoining fusion interface of both fragments and discarded. A second longitudinal cut was then made ∼0.5 mm from the shaved edge to isolate the recipient and donor boundary samples. Boundary samples therefore represented tissue immediately adjacent to, but not directly on, the fusion interface. Following boundary-sample removal, an additional longitudinal cut was made ∼5 mm from the newly exposed margin to obtain the donor and recipient far samples, which represented tissue distal to the fusion boundary.

To minimise cross-contamination, cutting tools were regularly cleaned between sampling by sequential dipping in ethanol, rinsing in freshwater and 0.2 µm FSW, and drying before reuse. Tissue sections from sample types 2a/b and 3a/b were trimmed into three ∼match-head-sized pieces. For sample types 1, 4, and 5, two ∼match-head-sized pieces were taken from the top and side surfaces of each coral fragment. All tissue pieces were rinsed in 0.2 µm FSW and preserved in absolute ethanol for subsequent DNA extraction. For sample types 2a/b and 3a/b, DNA extracted separately from the three tissue pieces was pooled prior to PCR amplification and ITS2 sequencing.

### DNA extraction and ITS2 sequencing

Details on DNA extraction, PCR amplification and library preparation are provided in Supplementary Information. Sequencing was conducted on an Illumina NextSeq v3 platform (2×300 bp) at the Walter and Eliza Hall Institute, Melbourne. Demultiplexed raw sequences were submitted to SymPortal for ITS2 profiling (Supplementary Information). The presence and abundance of recurring ITS2 sequences across samples are identified by SymPortal as ‘Internal Transcribed Spacer 2 (ITS2) type profiles’, which are representative of putative Symbiodiniaceae taxa. Each of the sequences in the recurring set (the profile) is referred to as a defining intragenomic variant (DIV). Sequencing data were downloaded from SymPortal and analysed in R version 4.2.0 (R Core Team, 2022). The ITS2 profile of the SS8 culture is C1-C1b-C42.2-C1bh-C1c-C1br-C1cb-C1ge-C1w-C3ju-C72k-C1al-C42ca-C3sa-C42ao-C1jx-C1cu [49]. Due to the relatively low abundance of SS8 in experimental corals, the presence of SS8 was primarily inferred from dominant SS8-associated DIVs (e.g., C1-C1b-C42.2). These DIVs provided sufficient resolution to assess putative SS8 acquisition in *G. fascicularis*. In contrast, ITS2 data from *P. columna* were excluded from strain-level symbiont analyses because high sequence similarity between SS8 and homologous *Cladocopium* spp. associated with this host precluded reliable discrimination of SS8. Accordingly, ITS2-based evidence for putative SS8 acquisition is restricted to *G. fascicularis*.

### Statistical analysis

All statistical analyses were conducted in R (v4.3.1). For both *P. columna* and *G. fascicularis*, a principal component analysis (PCA) was first performed on the RGB reflectance values extracted from linear transects across coral tissue. PC1 accounted for over 92% of the total variance in both species and was therefore used as the response variable in downstream analyses of reflectance. Changes in reflectance PC1 scores and *F_v_*/*F_m_* over time and among experimental categories were assessed using linear models and linear mixed-effects models. Paired *t-*tests were used to compare pre- and post-bleaching samples. For post-bleaching timepoints, model structure was selected using Akaike Information Criterion (AIC) comparisons. Genotype was included as a fixed effect, while fusion pairing was included as a random effect. Pairwise comparisons were conducted using estimated marginal means with Tukey adjustment.

## Results

### Fusion outcomes

Successful fusion was achieved in all donor–recipient pairings: 24 out of 24 in *Galaxea fascicularis* and 28 out of 28 in *Psammocora columna*. Fusion onset was rapid, with initial tissue integration observed as early as day thee in *G. fascicularis* and day nine in *P. columna* (Figs. 2A, B; Figs. 3A, B respectively; see Supplementary Information Table S1 for the full list of fusion onset times). While initial tissue contact occurred quickly, complete integration along the tissue boundary progressed more gradually. This slower progression appeared to reflect minor misalignments between donor and recipient margins, which required additional tissue growth to bridge remaining gaps.

**Fig 2.**
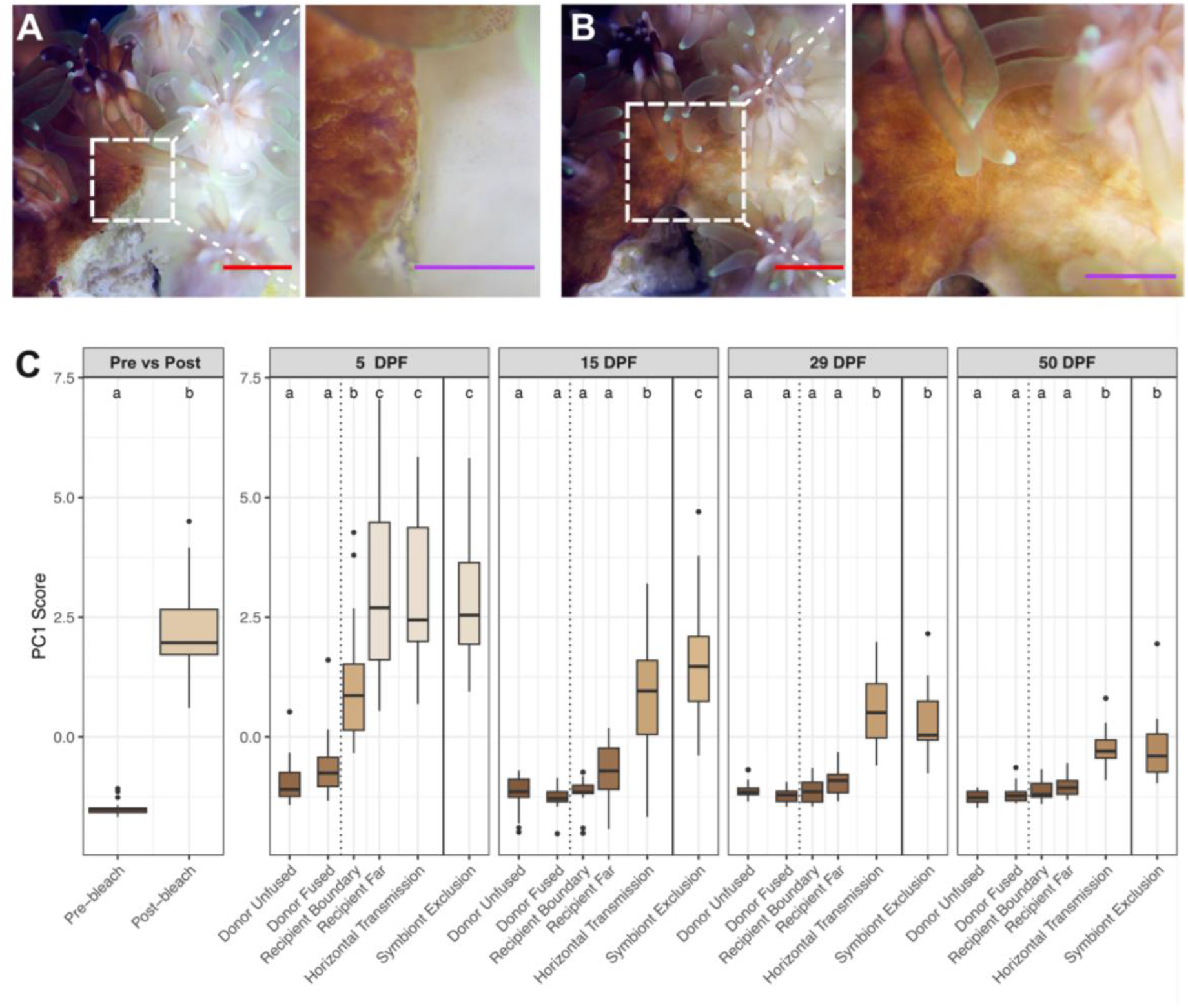
(A) *Galaxea fascicularis* donor (pigmented) and recipient (bleached) fragments at 0 DPF (days post-fusion setup). (B) The same donor-recipient pair at 6 DPF. Insets show magnified view of fusion line at both timepoints with clear tissue fusion observed at 6 DPF. Scale bars represent 2.5 mm (red) and 1 mm (purple). (C) Reflectance-based principal component analysis (PCA) scores (PC1) for *G. fascicularis* across timepoints and experimental categories. Each box represents the distribution of PC1 scores within a category at a given timepoint. The leftmost plots (Pre vs Post) compare coral reflectance profiles before and after chemical bleaching using a paired *t-*test. Experimental timepoints post-fusion track changes in PC1 scores over time across five experimental categories: unfused donor, fused donor, recipient boundary, recipient far, horizontal transmission, and symbiont exclusion. The colour scale reflects the mean PC1 score for each category, bounded by the minimum and maximum values observed across all timepoints. Category differences at each timepoint were assessed using post hoc pairwise contrasts from the fitted model. Different letters denote statistically significant groupings (Tukey-adjusted *P* < 0.05).

**Fig 3.**
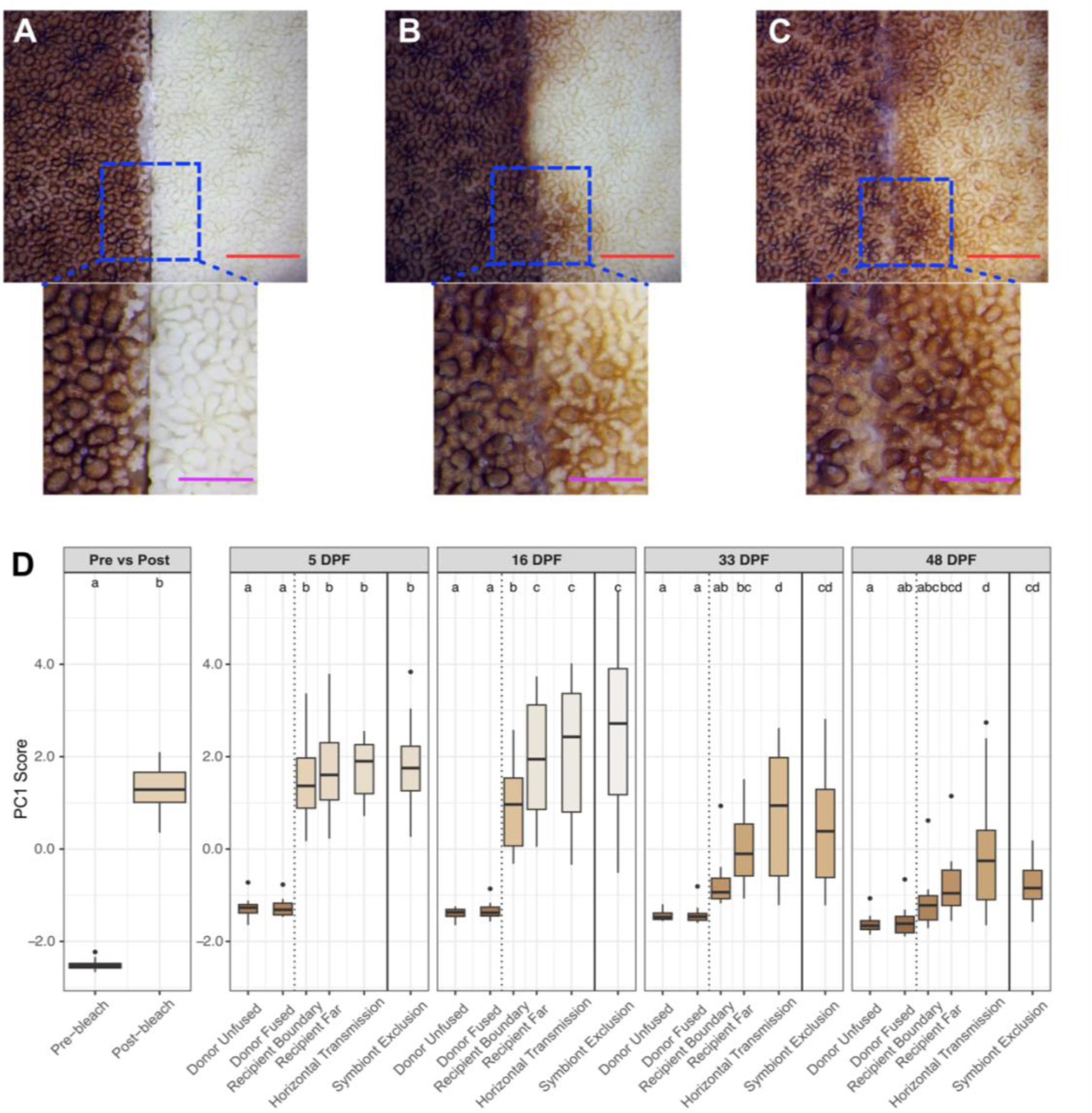
(A) *Psammocora columna* donor (pigmented) and recipient (bleached) fragments at 0 DPF (days post-fusion setup). (B) The same donor–recipient pair at 13 DPF and (C) 36 DPF (these images do not align temporally with reflectance measurements). Insets in both panels show magnified views of the fusion line revealing clear tissue fusion by 13 DPF. Scale bars represent 5 mm (red) and 2.5 mm (purple). (D) Reflectance-based principal component analysis (PCA) scores (PC1) for *P. columna* across timepoints and experimental categories. Each box represents the distribution of PC1 scores within a category at a given timepoint. The leftmost plots (Pre vs Post) compare coral reflectance profiles before and after chemical bleaching using a paired *t*-test. Experimental timepoints post-fusion track changes in PC1 scores over time across the five experimental categories: unfused donor (UD), fused donor (FD), recipient boundary, recipient far, and horizontal transmission (HT) and symbiont-exclusion (EC) controls. The colour scale reflects the mean PC1 score for each category, bounded by the minimum and maximum values observed across all timepoints. Category differences at each timepoint were assessed using post hoc pairwise contrasts from the fitted model. Different letters denote statistically significant groupings (Tukey-adjusted *P* < 0.05).

### Reflectance-based principal component analysis (PCA)

#### Galaxea fascicularis

A strong bleaching-associated shift in spectral profiles was observed in *G. fascicularis* following chemical treatment, with mean PC1 scores increasing significantly from −1.49 ± 0.15 (Pre-bleach) when fragments were fully pigmented to 2.20 ± 0.93 when tissue was visibly bleached (Post-bleach) (*P* < 0.05, paired *t*-test; Fig. 2C). Lower PC1 scores therefore corresponded to greater pigmentation, while higher PC1 scores corresponded to bleached tissue. Following bleaching confirmation, recipient and donor fragments were assigned to their respective treatments, and reflectance was tracked over time. Linear mixed-effects modelling revealed a significant treatment-by-timepoint interaction (*P* < 0.001), indicating divergent PC1 trajectories among experimental groups (Fig. 2C).

Both FD and UD tissues exhibited relatively consistent low and stable PC1 scores throughout the experiment, with no significant differences detected between treatments at any timepoint (P > 0.74). In contrast, all recipient treatments exhibited significantly elevated PC1 scores relative to donors at 5 days post-fusion setup (DPF). At this timepoint, recipient tissue nearer the boundary (recipient boundary) showed moderately elevated scores (1.12 ± 1.33) which were significantly higher than both FD and UD (P < 0.0001), but lower than those of recipient far, HT, and EC (P < 0.01 for all). By 15 DPF, recipient boundary scores declined sharply to −1.17 ± 0.33 and became statistically indistinguishable from both FD and UD (P > 0.99), a similarity that persisted through the end of the experiment. Recipient far tissue followed a similar trajectory, with PC1 scores dropping from 3.04 ± 1.99 at 5 DPF to −0.75 ± 0.65 at 15 DPF—becoming significantly lower than HT and EC (P < 0.0001 for both) and statistically similar to FD, UD (P > 0.1), and recipient boundary tissue (P = 0.56). This convergence was maintained at 29 and 50 DPF.

By 15 DPF, PC1 scores had declined in both HT and EC treatments (HT: 0.72 ± 1.26; EC: 1.55 ± 1.34), with a significantly greater reduction observed in HT compared to EC (*P* = 0.0076). Scores continued to decline at 29 DPF (HT: 0.52 ± 0.84; EC: 0.30 ± 0.74), by which point the difference between treatments was no longer statistically significant, although both remained elevated relative to all donor and recipient fusion zones. By 50 DPF, PC1 scores had further decreased to −0.27 ± 0.42 (HT) and −0.28 ± 0.65 (EC), yet both treatments were still significantly higher than donor and recipient fusion zones.

#### Psammocora columna

Following chemical bleaching, the mean PC1 scores of *P. columna* increased significantly from −2.51 ± 0.11 (Pre-bleach) to 1.31 ± 0.48 (Post-bleach) (*P* < 0.05, paired *t*-test; Fig. 3D). Linear mixed-effects modelling identified a significant treatment-by-timepoint interaction (*P* < 0.001), indicating divergent PC1 trajectories among experimental groups over time (Fig. 3D).

Across all timepoints, FD and UD treatments exhibited the lowest PC1 scores with no significant differences detected between the two donor treatments at any timepoint. In contrast, all recipient treatments displayed significantly elevated PC1 scores relative to donors at 5 DPF. No significant differences were detected among recipient treatments (*P* > 0.05) at this timepoint. At 16 DPF, although PC1 scores in recipient boundary tissue had declined (−34.7%), they remained significantly higher than donor treatments, positioning it as an intermediate between donor and other recipient treatments. From 33 DPF onwards, recipient boundary PC1 scores continued to decline (−0.76 ± 0.52 at 33 DPF; −1.16 ± 0.55 at 48 DPF), and became statistically indistinguishable from both FD and UD treatments (*P* > 0.05). Recipient far tissue followed a similar but more attenuated trajectory than recipient boundary with scores declining from 1.69 ± 0.94 at 5 DPF to −0.81 ± 0.68 at 48 DPF (-148%). At 16 DPF, recipient far tissue exhibited significantly higher PC1 scores than recipient boundary (2.00 ± 0.20 vs. 0.94 ± 0.20; *P* < 0.01) but remained lower than HT and EC treatments (*P* < 0.05).

By 33 DPF, recipient far tissue became statistically indistinguishable from both recipient boundary tissue (−0.07 ± 0.69 vs −0.76 ± 0.52; *P* = 0.08) and EC treatments (0.39 ± 1.22; *P* = 0.16); however, it remained significantly lower than HT (0.76 ± 1.42; P < 0.05). At 48 DPF, recipient far PC1 scores (−0.81 ± 0.68) were comparable with FD (−1.58 ± 0.31; *P* = 0.12), HT (−0.14 ± 1.33; *P* = 0.16), and EC treatments (−0.77 ± 0.49; *P* = 0.95), but remained significantly lower than UD corals (−1.63 ± 0.19; *P* = 0.03).

HT and EC corals exhibited the highest PC1 scores throughout the experiment. PC1 scores did not differ significantly between these treatments at any timepoint (*P* > 0.05). By 48 DPF, both treatments remained significantly higher than UD and FD corals (*P* < 0.05), but were no longer distinguishable from recipient far tissue. By 48 DPF, the EC treatment was also statistically indistinguishable from recipient boundary tissue.

### PSII maximum quantum yield (F_v_/F_m_)

The maximum quantum yield of photosystem II (*F_v_*/*F_m_*) declined significantly in both *G. fascicularis* and *P. columna* fragments following chemical bleaching (*P* < 0.0001, paired *t*-test). In *G. fascicularis*, *F_v_*/*F_m_* declined by 30.5%, from 0.456 ± 0.036 to 0.317 ± 0.037, while in *P. columna*, values decreased by 57.5%, from 0.478 ± 0.074 to 0.203 ± 0.055 (Fig. 4A, B). Thus, the proportional decline in *F_v_*/*F_m_* following chemical bleaching was almost twice as large in *P. columna* as in *G. fascicularis*. Following chemical bleaching, recipient and donor fragments were assigned to their respective treatments, and *F_v_*/*F_m_* was tracked over time. Linear mixed-effects modelling revealed a significant treatment-by-timepoint interaction for both species (*P* < 0.001), indicating distinct recovery trajectories across experimental groups (Fig. 4A, B).

**Fig 4.**
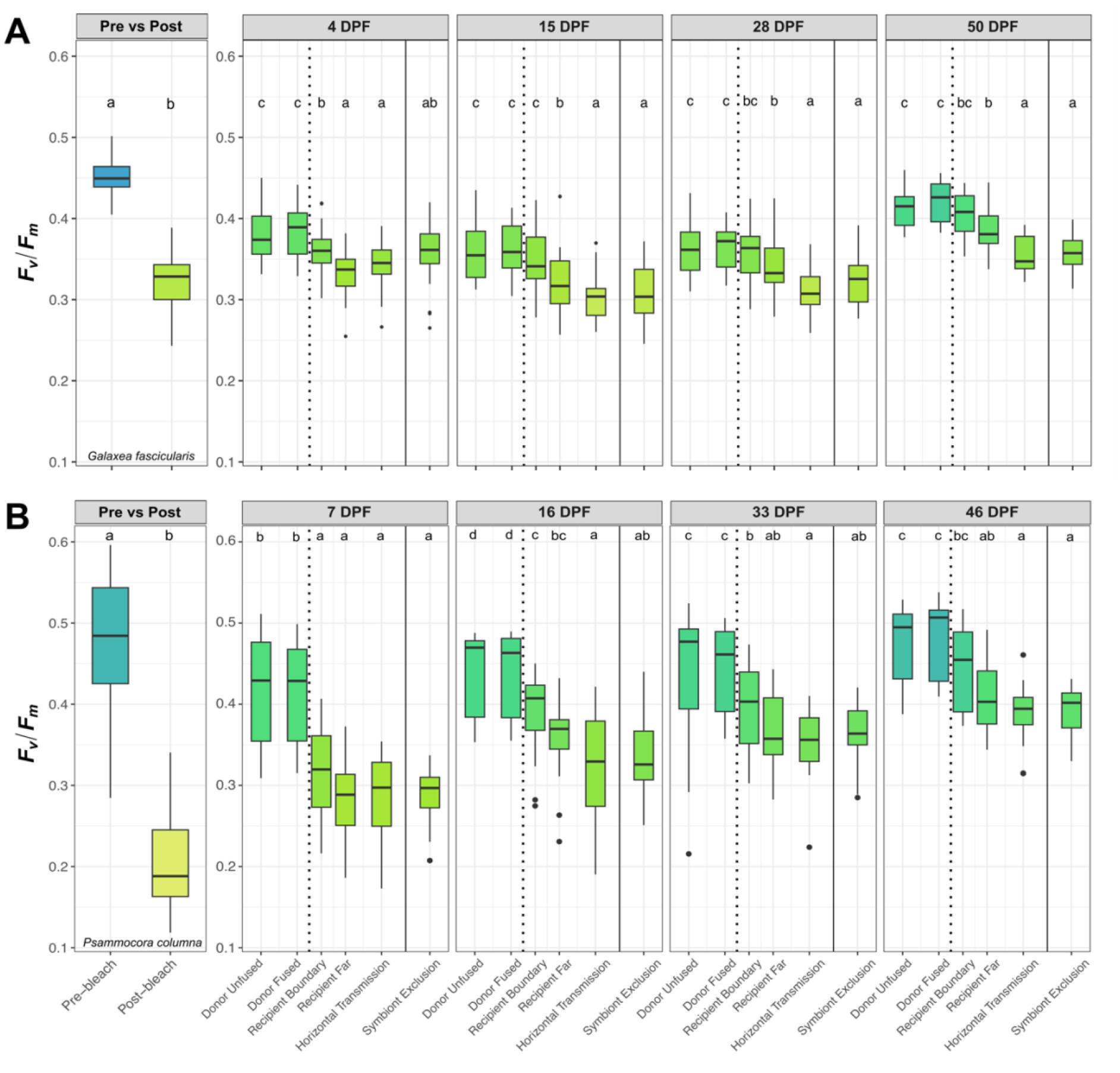
Maximum quantum yield of photosystem II (*F_v_*/*F_m_*) in (A) *Galaxea fascicularis* and (B) *Psammocora columna* across timepoints and experimental categories. Each boxplot displays the distribution of *F_v_*/*F_m_* values for a given category at each timepoint. The leftmost plots (Pre vs Post) compare *F_v_*/*F_m_* values before and after chemical bleaching using a paired *t*-test. All other timepoints represent post-bleaching recovery, with differences between categories assessed using linear mixed-effects models with fusion pairing specified as a random effect. The colour scale reflects the mean *F_v_*/*F_m_* value for each category, bounded by the minimum and maximum values observed across all timepoints. Different letters indicate statistically distinct groups based on Tukey-adjusted comparisons (*P* < 0.05).

In *G. fascicularis*, both FD and UD treatments had the highest *F_v_*/*F_m_* values across all timepoints (0.362 to 0.422), with no significant differences detected between them. *F_v_*/*F_m_* generally increased over time in both recipient boundary and recipient far tissues, although all treatments showed a slight decline between 4 and 15 DPF. Recipient boundary values rose from 0.362 ± 0.031 at 4 DPF to 0.404 ± 0.025 by the end of the experiment at 50 DPF (+11.6%), and recipient far increased from 0.333 ± 0.031 to 0.387 ± 0.026 (+16.2%). At 4 DPF, recipient boundary tissue exhibited significantly higher *F_v_*/*F_m_* values than both recipient far and HT treatments but did not differ from EC corals. By 15 DPF, *F_v_*/*F_m_* of recipient boundary tissue was significantly higher than all other recipient treatments and statistically indistinguishable from both donor treatments, a pattern that persisted across subsequent timepoints. By 15 DPF, recipient far shifted towards an intermediate position relative to the other recipient treatments. It became statistically indistinguishable from recipient boundary tissue from 28 DPF onward, while remaining significantly lower than both donor treatments throughout the experiment.

HT and EC treatments consistently exhibited the lowest *F_v_*/*F_m_* values throughout the experiment, except at 4 DPF when recipient far recorded the lowest value (0.333 ± 0.031). In HT corals, *F_v_*/*F_m_* increased from a minimum of 0.302 ± 0.029 at 15 DPF to 0.355 ± 0.023 by 50 DPF (+17.5%). A similar trend was observed in EC corals, rising from 0.309 ± 0.037 at 15 DPF to 0.356 ± 0.023 at 50 DPF (+15.2%). No significant differences were detected between HT and EC treatments at any timepoint. Across all timepoints, both treatments had significantly lower *F_v_*/*F_m_* values than recipient boundary tissue, except at 4 DPF when EC values did not differ significantly from those of recipient boundary.

In *P. columna*, FD and UD treatments consistently exhibited the highest *F_v_*/*F_m_* values (0.414– 0.477) across all timepoints. No differences were detected between these two treatments throughout the experiment. Among recipient treatments, recipient boundary fragments recovered to the highest absolute *F_v_*/*F_m_* value, increasing from 0.314 ± 0.054 at 7 DPF to 0.442 ± 0.052 at 46 DPF (+40.8%). At 16 and 46 DPF, the *F_v_*/*F_m_* of recipient boundary tissue was significantly higher than both HT and EC controls; at 33 DPF, it was significantly higher than HT but not EC. By 46 DPF, recipient boundary tissue was no longer significantly different from either donor treatment. Recipient far tissue followed a similar pattern, increasing from 0.285 ± 0.046 to 0.406 ± 0.044 (+42.5%). Although consistently lower, recipient far tissue did not differ significantly from recipient boundary tissue at any timepoint. The *F_v_*/*F_m_* of recipient far tissue also remained significantly lower than both donor treatments throughout the experiment.

The HT and EC treatments consistently exhibited the lowest *F_v_*/*F_m_* values, except at 7 DPF when recipient far tissue recorded the lowest value (0.285 ± 0.046). Across the experiment, *F_v_*/*F_m_* in HT fragments increased by 35.8% (from 0.286 ± 0.053 to 0.389 ± 0.034), and in EC fragments by 35.8% (from 0.290 ± 0.037 to 0.392 ± 0.031). No significant differences were observed between HT and EC treatments at any timepoint.

### Galaxea fascicularis ITS2 Symbiodiniaceae community profiles

Prior to the start of the fusion experiment (0 DPF; ∼50 days post-SS8 inoculation), SS8 was detected in 3 of 19 *G. fascicularis* donor fragments, at low relative abundances of 0.68%, 0.84%, and 1.32%, with *Durusdinium* remaining the dominant symbiont (>98%; see Supplementary Information Fig S4). SS8 was not detected in any recipient fragments either before or after chemical bleaching (see Supplementary Information Fig S5). Bleaching substantially reduced the relative abundance of native *Cladocopium* in recipients: among the five of 19 fusion pairings that contained native *Cladocopium* prior to chemical bleaching, only three retained detectable levels post-bleaching, with average relative abundances decreasing from 13.08 ± 4.68% to 6.06 ± 5.32% (mean ± SE).

Although SS8 was detected in only a subset of donor fragments sampled at 0 DPF, it was detected more broadly in fused donor tissues by the final timepoint. Detection occurred in donor far (3/18), donor boundary (8/18), or both regions (7/18), with relative abundances in FDs ranging from 0.94% to 5.65% (mean ± SE: 2.21 ± 0.21%). No apparent differences in SS8 abundance were observed between donor boundary and donor far sampling regions (1.91 ± 0.23% vs. 2.41 ± 0.30%, respectively). SS8 was, however, detected more frequently in FD sampling regions (18/19) than in UD corals (11/18; one UD coral was excluded due to insufficient sequencing depth). Patterns of SS8 co-occurrence across donor, recipient, and HT categories are summarised in Supplementary Information Table S2.

Among fused recipients whose paired donor also had detectable SS8 at the final timepoint, SS8 was detected in 15 of 19 corals, occurring in recipient boundary tissue (2/15), recipient far tissue (1/15), or both (12/15). Relative abundances ranged from 0.75% to 3.14% (mean ± SE = 1.56 ± 0.10%). No apparent differences in SS8 abundance were observed between recipient boundary and recipient far sampling regions (1.59 ± 0.12% vs. 1.53 ± 0.16%, respectively). SS8 was also detected in 13 out of 18 HT fragments, where it occurred at slightly higher relative abundances than in fused-recipient tissue, ranging from 1.13% to 9.28% (mean ± SE = 3.54 ± 0.67%).

Across all treatments, only symbionts belonging to the genera *Cladocopium* and *Durusdinium* were detected, with *Durusdinium* remaining the dominant genus. Residual native *Cladocopium* was again detected in only a small subset of corals (4 of 19), spanning 11 distinct tissue sampling regions, with an average relative abundance of 12.63 ± 6.43%—slightly higher than the average observed in recipients immediately post-bleaching. No SS8 DIVs were detected in EC corals, suggesting no detectable cross-contamination prior to or during the experiment.

Across all samples, SS8 relative abundance was significantly influenced by treatment/sampling region (two-way ANOVA: *F*(5, 59) = 4.04, *P* = 0.0032), but not by genotype (*F*(2, 59) = 1.07, *P* = 0.35), with no interaction observed between treatment/sampling region and genotype (*F*(10, 59) = 0.37, *P* = 0.95). Post hoc comparisons indicated that SS8 abundance was significantly higher in HT samples compared to both recipient boundary and recipient far tissues (*P* = 0.0108 and 0.0085, respectively), while no other pairwise differences among treatment categories/sampling regions were significant.

## Discussion

This study tested whether adult isograft fusion between healthy “donor” corals hosting heat-evolved (HE) *Cladocopium proliferum* (strain SS8) and chemically bleached “recipients” could facilitate SS8 acquisition and post-bleaching recovery. Despite SS8 being present at low relative abundance (<1.32%) in *Galaxea fascicularis* donors at the start of the experiment, SS8 was detected in 15 of 19 fused recipients whose paired donor also had detectable SS8 by the final timepoint. However, SS8 was also detected in unfused recipients in the horizontal transmission (HT) control treatment, where its prevalence was slightly but significantly higher, consistent with previous evidence that corals can acquire expelled symbiont cells from the surrounding water column [31]. SS8 detection and relative abundance also showed no consistent spatial gradient between tissues sampled near the fusion interface and those sampled farther from it. This contrasts with the clear spatial pattern in physiological recovery, whereby tissue pigmentation and photochemical efficiency improved more rapidly near the fusion interface in recipients of both *G. fascicularis* and *Psammocora columna*. Together, these findings indicate that adult coral tissue fusion can accelerate spatially localised physiological recovery and coincide with detectable SS8 in recipient tissues. The absence of a corresponding spatial pattern in SS8 abundance, however, indicates that the localised physiological response cannot be attributed solely to the accumulation of SS8 transferred directly across the fusion interface. The results therefore do not fully disentangle direct tissue-mediated transfer from water-column acquisition, proliferation of residual recipient symbionts, or other physiological benefits associated with tissue continuity between fused fragments.

### Fusion between isografts

Of the 52 successful fusion pairings across both species, no instances of incompatible fusion reactions—i.e., non-fusion or tissue necrosis at the interface—were observed throughout the ∼50-day experiment. This outcome was expected, as all fragments in a pair were clonally derived from the same original “mother” fragment and were therefore genetically identical isografts. Indeed, microfragmentation and fusion of clonal fragments is a well-established aquaculture technique commonly used to accelerate coral growth [37, 40]. Although chemical bleaching can impose substantial physiological stress on coral hosts and their symbionts, fusion success was not compromised under the conditions used here.

### Fusion to a healthy donor accelerates bleaching recovery

Following tissue fusion, the gastrovascular systems (GVS) of the two colonies are expected to integrate [44], potentially facilitating exchange of dissolved and particulate materials, including organic and inorganic nutrients and, in some cases, freely circulating microalgal symbionts [44, 45]. In this study, fused recipients of both species showed visual and physiological patterns in line with localised symbiont recovery near the fusion interface. Macroscopic reflectance imaging revealed greater pigmentation in fused recipients than in unfused controls, with recovery strongest near the fusion boundary and declining with distance from the interface (Figs. 2, 3). Symbiont photochemical efficiency also improved in a similar spatial pattern, suggesting coordinated recovery of both symbiont-associated pigmentation and photochemical performance (Figs. 4A, B). The magnitude of photochemical recovery nevertheless differed between species. Recipient boundary and recipient far tissues increased by 40.9% and 42.6%, respectively, in *P. columna*, compared with 11.6% and 16.2% in *G. fascicularis*. The larger proportional increases in *P. columna* likely partly reflect its lower *F_v_/F_m_* immediately post-bleaching, consistent with greater sensitivity to the chemical bleaching treatment. However, interpretation of the mechanistic basis of these patterns is complicated by the nature of chemical bleaching, which removes most—but not all—symbionts from coral host tissues [49]. Consequently, it remains unclear whether recovery reflects the enhanced restoration of photochemical function and or proliferation of residual recipient symbionts, uptake of expelled donor cells, direct transfer through the fused tissue interface, or a combination of these processes. These possibilities are considered below (see Fig. 6 for a conceptual overview).

**Fig 5.**
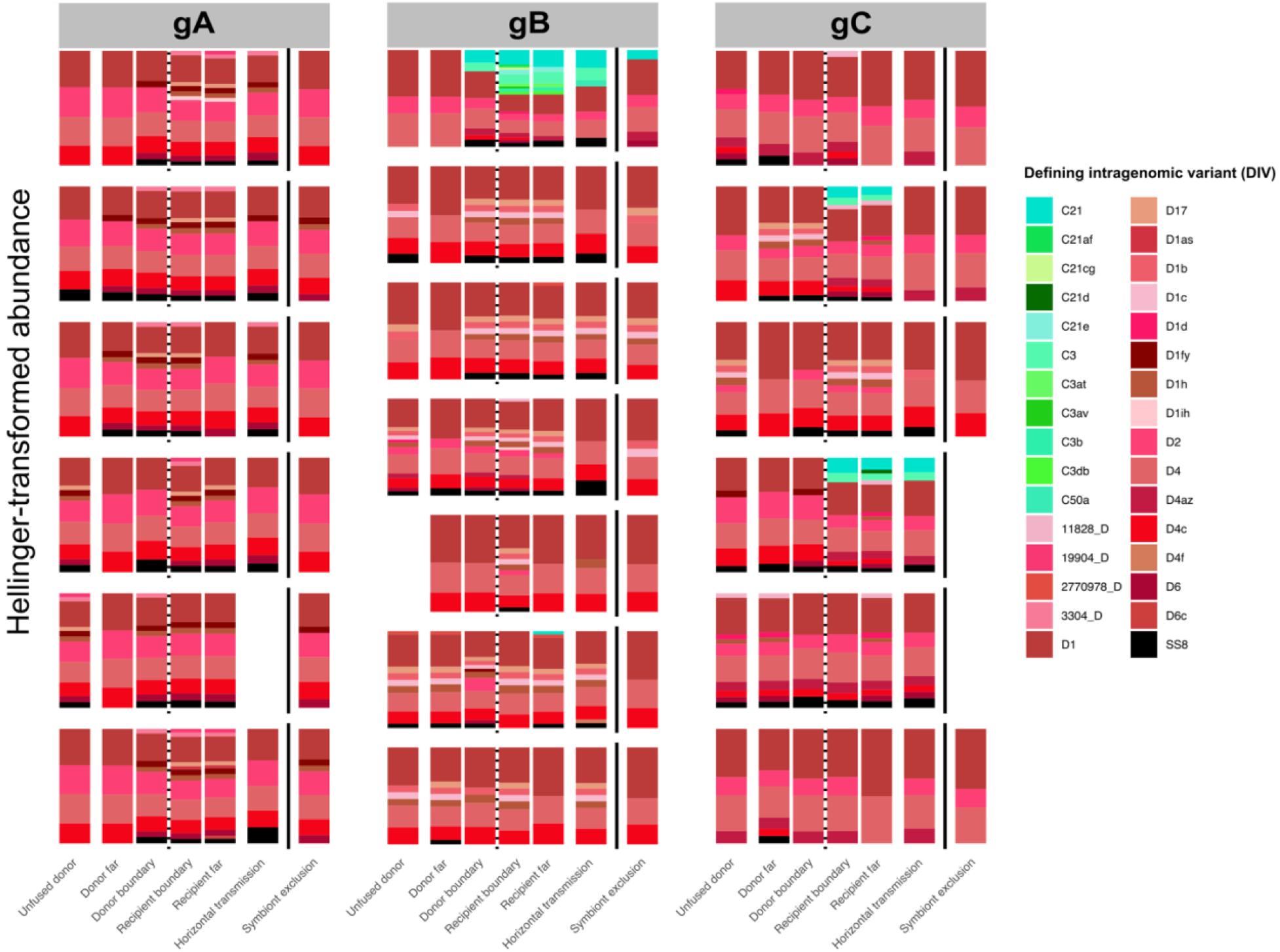
*Galaxea fascicularis* ITS2 Symbiodiniaceae community composition across experimental treatments. Bar plots show ITS2-derived community profiles at the final timepoint (51 DPF) for three *G. fascicularis* genotypes (gA, gB, and gC). Each plot represents an individual fragment pairing, with bars indicating the proportional abundance of dominant intragenomic variants (DIVs), coloured by genus or identified as SS8 (black). Treatments are faceted as: unfused donor (UD), donor far, donor boundary, recipient boundary, recipient far, and horizontal transmission (HT) and symbiont-exclusion (EC) controls. Empty or missing bars indicate samples that did not pass quality control.

**Fig 6.**
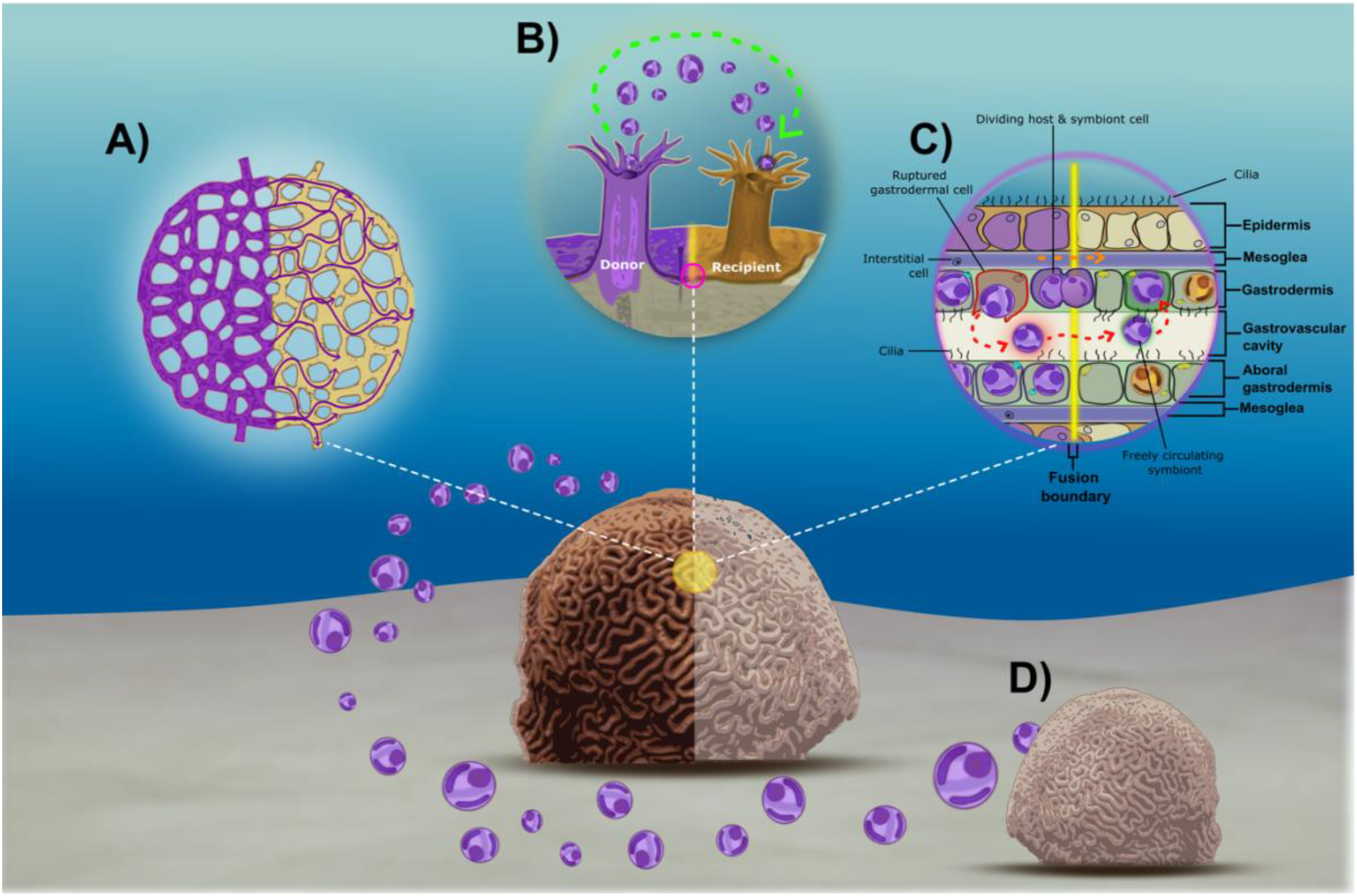
Conceptual diagram illustrating potential mechanisms of heat-evolved (HE) symbiont and nutrient transfer across coral fragments following tissue fusion. (A) Widespread diffusion of nutrients and metabolites (either passively or via active transport) from donor (purple) to recipient (yellow) across the fusion line; (B) localised expulsion of HE symbionts from donor polyp and subsequent acquisition by recipient at or near the fusion line; (C) integration of the gastrovascular cavities following fusion, potentially allowing donor-derived symbionts that have been released to circulate through the shared gastrovascular cavity and be acquired by recipient gastrodermal cells (red arrows). Also shown is the potential movement or proliferation of symbiont-bearing host cells across the fusion line (orange arrows); (D) expulsion of HE symbionts into the water-column for potential acquisition by neighbouring bleached corals.

### Fusion-mediated nutrient diffusion and recovery of residual symbionts

One possible explanation for the localised recovery observed in fused recipients is that fusion enabled the transfer of donor-derived metabolites into bleached recipient tissue, supporting host recovery and the proliferation of residual symbionts (Fig. 6A). Indeed, previous work in the soft coral *Pseudopterogorgia fulvum fulvum* has shown that, following fusion, ¹⁴C-labelled photosynthates can move into the recipient colony and reach tissues up to 390 mm from the fusion line within 24 hours [44]. Although this study was conducted on a soft coral (Alcyonacea), similar resource-sharing dynamics may occur in scleractinian corals, including *G. fascicularis*, which possesses a mesh-like canal system thought to support uniform nutrient transport among polyps [51]. Transferred metabolites could include organic carbon and nitrogenous compounds that support host metabolism and Symbiodiniaceae growth, given that ammonium and other host-derived nutrients are readily assimilated by Symbiodiniaceae and can support protein synthesis and cell division [52–54]. However, as nutrient fluxes were not measured here, metabolite transfer remains a plausible but untested contributor to the recovery patterns observed here.

### Localised polyp-scale transfer of expelled symbionts from donor to recipient

A second, non-mutually exclusive explanation is that symbionts expelled from donor polyps were locally acquired by adjacent recipient tissue near the fusion interface (Fig. 6B). This explanation is consistent with the detection of SS8 in horizontal transmission controls, which indicates that expelled or otherwise released SS8 cells were available in the surrounding water column for acquisition by bleached recipients [31]. In fused pairs, the close proximity of donor and recipient polyps may have increased the local density of released symbionts near the fusion boundary, providing a spatially localised source of cells for uptake. Local retention of released symbionts could also be influenced by coral surface flows. The coral surface is surrounded by a diffusive boundary layer (DBL), which can be modified by epidermal ciliary beating to enhance solute transport and retain small particles in the mucus layer [55–58]. For example, ciliary flow in *Porites lutea* has been shown to entrain suspended particles (∼5 μm), concentrating them within the overlying mucus and increasing retention 3.4-fold [56]. Similar near-surface flow processes could therefore increase the residence time of expelled symbionts near recipient tissues, particularly where donor and recipient polyps are in direct contact. Interestingly, however, SS8 relative abundance was slightly higher in HT recipients than in fused recipient tissues, indicating no clear enhancement of SS8 abundance associated with proximity to donor polyps. Nonetheless, because expelled-cell viability, uptake, and functional contribution were not directly measured, this pathway remains a plausible but unresolved contributor to localised recovery.

### Fused donor to recipient symbiont migration

A third, non-mutually exclusive explanation for the earlier and more localised recovery observed in fused recipients is that tissue fusion may have permitted symbionts to move directly through newly connected gastrovascular or gastrodermal tissues (Fig. 6C; red arrow). Freely circulating Symbiodiniaceae cells have been directly observed within the coelenteron of reef-building corals, including *Acropora cervicornis* [45], as well as in several octocoral species, where such movement has been proposed to contribute to bleaching recovery [59]. Under this scenario, donor-derived symbionts released into a shared gastrovascular cavity could move into recipient regions where they may be captured by ciliated endodermal cells— a process consistent with previously described mechanisms of symbiont uptake in corals [60].

In addition to movement via the gastrovascular cavity, localised recovery may reflect redistribution of symbionts through newly connected gastrodermal tissue, potentially involving movement or proliferation of symbiont-bearing host cells across the fusion interface (Fig. 6C; orange arrow). This possibility is supported by observations from Bockel and Rinkevich [61], who documented rapid migration of endosymbiotic dinoflagellates from densely populated tissues into newly developed, symbiont-free tissue in Red Sea pocilloporid corals. However, the precise cellular mechanisms underlying rapid symbiont movement through host tissues remain largely unknown.

It is important to note that the above mechanisms are not mutually exclusive and may operate sequentially or synergistically. Donor-derived metabolites could enhance host condition and support the proliferation of residual symbionts, while ciliary modification of the DBL could increase the local retention and uptake of expelled cells near the fusion interface. Gastrovascular integration could provide an additional internal route of transport following tissue fusion. However, because nutrient fluxes, near-surface hydrodynamics, and symbiont movement were not measured directly, their relative and interactive contributions cannot be distinguished from the present data.

Although this study primarily examined the impact of fusion on the bleached recipient, fusion could also be expected to affect the donor if nutrients or symbionts are shared with an attached, resource-limited partner. However, both reflectance and photochemical efficiency were indistinguishable between fused and unfused donor corals, suggesting no detectable physiological cost to the donor under the experimental conditions and timeframe here. This apparent asymmetry may reflect a broader principle of modularity in colonial organisms, in which the ability to integrate additional units at minimal physiological cost represents a key evolutionary advantage of the coral colonial lifestyle.

### Future directions

Several important questions remain before the broader ecological relevance of fusion-mediated recovery and symbiont acquisition can be established. First, future work should determine whether similar patterns occur in allografts between genetically distinct conspecifics, or in xenografts between closely related species. Determining this will be critical for assessing the ecological relevance of this method. Although allograft and xenograft pairings often lead to tissue rejection or necrosis, Nozawa and Loya [43] showed that juvenile *Seriatopora* allografts and xenografts can exhibit signs of early-stage fusion that only later transition into tissue separation or necrotic disconnection as the allorecognition system matures. Whether a comparable window of transient fusion exists in adult corals with more developed allorecognition systems remains unknown; however, even short periods of tissue integration may be sufficient to permit resource exchange or symbiont movement across the fusion boundary.

Future work should also determine whether low-abundance SS8 contributes measurably to coral performance. In the present study, SS8 was detected in fused *G. fascicularis* recipients, but its relative abundance was generally low and its functional contribution to photochemical recovery could not be separated from residual recipient symbiont proliferation or donor-derived nutritional support. Controlled dose–response experiments, longer-term monitoring, and subsequent heat-stress assays will be needed to test whether acquired HE symbionts can meaningfully alter post-bleaching recovery or thermal tolerance.

A major challenge in studies of symbiont transmission and establishment will be disentangling the mechanisms by which introduced symbionts arrive and persist within recipient tissues. Specifically, future studies will need to control for potential horizontal acquisition from the surrounding water column in order to distinguish this pathway from direct symbiont transfer across the fusion interface. One potential approach would be to develop an agarose-based system designed to restrict the dispersal of expelled symbionts and reduce their transport by coral-generated ciliary currents. By limiting water movement around the fragments while preserving direct tissue contact at the fusion interface, such a system could provide a more stringent test of whether symbionts move directly between fused partners. Residual symbiont proliferation represents another possible explanation for recipient recovery and will also require further examination. Live-cell imaging at the fusion line, particularly where donor- and recipient-derived symbionts can be differentiated, could help resolve the real-time dynamics of symbiont movement, cellular division, and incorporation into recipient tissues. In addition to symbiont acquisition and proliferation, nutrient exchange between fused partners may also contribute to recovery. Stable isotope labelling of donor colonies, such as with ¹³C-bicarbonate or ¹⁵N-nitrate, could be used to assess the direction, extent, and spatial distribution of nutrient flow across the fusion interface. Although this approach would not resolve symbiont identity, it would help clarify whether localised metabolic enrichment contributes to recipient recovery.

A further limitation of the present study is the lack of tank replication across SS8-exposed and symbiont-exclusion controls. Because each group was housed in a single tank, SS8-related effects cannot be fully separated from other tank-specific differences in environmental conditions or husbandry. Replication of both groups across multiple independent tanks would therefore be required to attribute observed differences more confidently to SS8 exposure rather than tank-level variation.

### Conclusion

This study provides a controlled first test of adult isograft fusion as a potential pathway for localised recovery and acquisition of experimentally evolved heat-tolerant Symbiodiniaceae. Fusion with SS8-associated donor fragments accelerated pigmentation recovery and enhanced photochemical efficiency in chemically bleached recipients, with recovery strongest near the fusion interface. In *G. fascicularis*, SS8 was detected in most fused recipients, consistent with HE symbiont acquisition, although its presence in horizontal transmission controls indicates that water-column uptake also contributed. These findings suggest that adult coral tissue fusion can support localised post-bleaching recovery and may contribute to symbiont movement between coral fragments. However, the relative contributions of direct tissue-mediated transfer, water-column acquisition, residual symbiont proliferation, and donor-derived nutrient exchange remain unresolved.

## Acknowledgements

The authors gratefully thank Elizabeth Kaplan for her invaluable support in the laboratory and her meticulous attention to detail. WYC acknowledges the Australian Research Council Discovery Early Career Researcher Award (DE240100317) and the Westpac Research Fellowship. This research was supported by the Australian Research Council Laureate Fellowship to MJHvO (FL180100036), Allen Family Philanthropies, the Reef Restoration and Adaptation Program—funded by the partnership between the Australian Government’s Reef Trust and the Great Barrier Reef Foundation—and the University of Melbourne. MRN acknowledges a Royal Society Te Apārangi Marsden Fast Start grant (21-VUW-021).

## Data Availability Statement

Raw ITS2 Symbiodiniaceae sequence data will be deposited in GenBank prior to publication and made publicly available upon acceptance. Accession numbers will be included in the revised manuscript. The authors declare that there is no conflict of interest associated with this research.

## References

[1] G. Muller-Parker, C. F. D’Elia, and C. B. Cook, “Interactions Between Corals and Their Symbiotic Algae,” in Coral Reefs in the Anthropocene, C. Birkeland, Ed. Dordrecht: Springer Netherlands, 2015, pp. 99–116.

[2] L. Muscatine and V. Weis, “Productivity of Zooxanthellae and Biogeochemical Cycles,” in Primary Productivity and Biogeochemical Cycles in the Sea, P. G. Falkowski and A. D. Woodhead, Eds. Boston, MA: Springer US, 1992, pp. 257–271.

[3] S. K. Davy, D. Allemand, and V. M. Weis, “Cell biology of cnidarian-dinoflagellate symbiosis,” Microbiol. Mol. Biol. Rev., vol. 76, no. 2, pp. 229–261, 2012.

[4] F. Lipschultz and C. Cook, “Uptake and assimilation of ^15^N-ammonium by the symbiotic sea anemones *Bartholomea annulata* and *Aiptasia pallida*: conservation versus recycling of nitrogen,” Mar. Biol., vol. 140, no. 3, pp. 489–502, 2002.

[5] J. Wiedenmann, C. D’Angelo, M. L. Mardones et al., “Reef-building corals farm and feed on their photosynthetic symbionts,” Nature, vol. 620, no. 7976, pp. 1018–1024, 2023.

[6] L. Muscatine and J. W. Porter, “Reef corals: mutualistic symbioses adapted to nutrient-poor environments,” BioScience, vol. 27, no. 7, pp. 454–460, 1977.

[7] V. M. Weis, “Cellular mechanisms of Cnidarian bleaching: stress causes the collapse of symbiosis,” J. Exp. Biol., vol. 211, no. 19, pp. 3059–3066, 2008.

[8] O. Hoegh-Guldberg, “Climate change, coral bleaching and the future of the world’s coral reefs,” Mar. Freshw. Res., vol. 50, no. 8, pp. 839–866, 1999.

[9] R. Cunning and A. C. Baker, “Thermotolerant coral symbionts modulate heat stress-responsive genes in their hosts,” Mol. Ecol., vol. 29, no. 15, pp. 2940–2950, 2020.

[10] K. E. Turnham, M. D. Aschaffenburg, D. T. Pettay et al., “High physiological function for corals with thermally tolerant, host-adapted symbionts,” Proc. R. Soc. B Biol. Sci., vol. 290, no. 2003, p. 20231021, 2023.

[11] A. M. Palacio-Castro, T. B. Smith, V. Brandtneris et al., “Increased dominance of heat-tolerant symbionts creates resilient coral reefs in near-term ocean warming,” Proc. Natl. Acad. Sci., vol. 120, no. 8, p. e2202388120, 2023.

[12] T. C. LaJeunesse, D. C. Wham, D. T. Pettay et al., “Ecologically differentiated stress-tolerant endosymbionts in the dinoflagellate genus *Symbiodinium* (Dinophyceae) Clade D are different species,” Phycologia, vol. 53, no. 4, pp. 305–319, 2014.

[13] K. D. Hoadley, A. M. Lewis, D. C. Wham et al., “Host–symbiont combinations dictate the photo-physiological response of reef-building corals to thermal stress,” Sci. Rep., vol. 9, no. 1, p. 9985, 2019.

[14] R. Berkelmans and M. J. van Oppen, “The role of zooxanthellae in the thermal tolerance of corals: a ‘nugget of hope’ for coral reefs in an era of climate change,” Proc. R. Soc. B Biol. Sci., vol. 273, no. 1599, pp. 2305–2312, 2006.

[15] R. N. Silverstein, R. Cunning, and A. C. Baker, “Change in algal symbiont communities after bleaching, not prior heat exposure, increases heat tolerance of reef corals,” Global Change Biol., vol. 21, no. 1, pp. 236–249, 2015.

[16] A. F. Little, M. J. van Oppen, and B. L. Willis, “Flexibility in algal endosymbioses shapes growth in reef corals,” Science, vol. 304, no. 5676, pp. 1492–1494, 2004.

[17] N. E. Cantin, M. J. van Oppen, B. L. Willis et al., “Juvenile corals can acquire more carbon from high-performance algal symbionts,” Coral Reefs, vol. 28, pp. 405–414, 2009.

[18] R. Cunning, P. Gillette, T. Capo et al., “Growth tradeoffs associated with thermotolerant symbionts in the coral *Pocillopora damicornis* are lost in warmer oceans,” Coral Reefs, vol. 34, no. 1, pp. 155–160, 2015.

[19] S. B. Matsuda, M. L. Opalek, R. Ritson-Williams et al., “Symbiont-mediated tradeoffs between growth and heat tolerance are modulated by light and temperature in the coral Montipora capitata,” Coral Reefs, vol. 42, no. 6, pp. 1385–1394, 2023.

[20] D. W. Kemp, K. D. Hoadley, A. M. Lewis et al., “Thermotolerant coral–algal mutualisms maintain high rates of nutrient transfer while exposed to heat stress,” Proc. R. Soc. B Biol. Sci., vol. 290, no. 2007, p. 20231403, 2023.

[21] I. Yuyama and T. Higuchi, “Comparing the effects of symbiotic algae (Symbiodinium) clades C1 and D on early growth stages of Acropora tenuis,” PLoS One, vol. 9, no. 6, p. e98999, 2014.

[22] D. C. Claar, S. Starko, K. L. Tietjen et al., “Dynamic symbioses reveal pathways to coral survival through prolonged heatwaves,” Nat. Commun., vol. 11, no. 1, p. 6097, 2020.

[23] M. J. van Oppen, J. K. Oliver, H. M. Putnam et al., “Building coral reef resilience through assisted evolution,” Proc. Natl. Acad. Sci., vol. 112, no. 8, pp. 2307–2313, 2015.

[24] M. R. Nitschke, D. Abrego, C. E. Allen et al., “The use of experimentally evolved coral photosymbionts for reef restoration,” Trends Microbiol., vol. 32, no. 12, pp. 1241–1252, 2024.

[25] I. E. Huertas, M. Rouco, V. Lopez-Rodas et al., “Warming will affect phytoplankton differently: evidence through a mechanistic approach,” Proc. R. Soc. B Biol. Sci., vol. 278, no. 1724, pp. 3534–3543, 2011.

[26] L. J. Chakravarti, V. H. Beltran, and M. J. van Oppen, “Rapid thermal adaptation in photosymbionts of reef-building corals,” Global Change Biol., vol. 23, no. 11, pp. 4675–4688, 2017.

[27] L. J. Chakravarti and M. J. van Oppen, “Experimental evolution in coral photosymbionts as a tool to increase thermal tolerance,” Front. Mar. Sci., vol. 5, p. 227, 2018.

[28] W. Y. Chan, L. Meyers, D. Rudd et al., “Heat-evolved algal symbionts enhance bleaching tolerance of adult corals without trade-off against growth,” Global Change Biol., vol. 29, no. 24, pp. 6945–6968, 2023.

[29] K. M. Quigley, C. Alvarez-Roa, J.-B. Raina et al., “Heat-evolved microalgal symbionts increase thermal bleaching tolerance of coral juveniles without a trade-off against growth,” Coral Reefs, vol. 42, no. 6, pp. 1227–1232, 2023.

[30] P. Buerger, C. Alvarez-Roa, C. Coppin et al., “Heat-evolved microalgal symbionts increase coral bleaching tolerance,” Sci. Adv., vol. 6, no. 20, p. eaba2498, 2020.

[31] B. G. Johnston, M. R. Nitschke, W. Y. Chan et al., “Horizontal transmission of heat-evolved microalgal symbionts in adult corals,” ISME J., vol. 19, no. 1, p. wraf157, 2025.

[32] K.-O. Amar, N. E. Chadwick, and B. Rinkevich, “Coral kin aggregations exhibit mixed allogeneic reactions and enhanced fitness during early ontogeny,” BMC Evol. Biol., vol. 8, no. 1, p. 126, 2008.

[33] M. Hidaka, “Tissue compatibility between colonies and between newly settled larvae of *Pocillopora damicornis*,” Coral Reefs, vol. 4, no. 2, pp. 111–116, 1985.

[34] L. Jiang, X.-M. Lei, S. Liu et al., “Fused embryos and pre-metamorphic conjoined larvae in a broadcast spawning reef coral,” F1000Research, vol. 4, p. 44, 2015.

[35] C. A. Ligson, P. C. Cabaitan, and P. L. Harrison, “Survival and growth of coral recruits in varying group sizes,” J. Exp. Mar. Biol. Ecol., vol. 556, p. 151793, 2022.

[36] L. J. Raymundo and A. P. Maypa, “Getting bigger faster: mediation of size-specific mortality via fusion in juvenile coral transplants,” Ecol. Appl., vol. 14, no. 1, pp. 281–295, 2004.

[37] Z. H. Forsman, B. Rinkevich, and C. L. Hunter, “Investigating fragment size for culturing reef-building corals (*Porites lobata* and *P. compressa*) in *ex situ* nurseries,” Aquaculture, vol. 261, no. 1, pp. 89–97, 2006.

[38] M. Hidaka, K. Yurugi, S. Sunagawa et al., “Contact reactions between young colonies of the coral *Pocillopora damicornis*,” Coral Reefs, vol. 16, no. 1, pp. 13–20, 1997.

[39] B. Rinkevich and Y. Loya, “Oriented translocation of energy in grafted reef corals,” Coral Reefs, vol. 1, no. 4, pp. 243–247, 1983.

[40] C. A. Page, E. M. Muller, and D. E. Vaughan, “Microfragmenting for the successful restoration of slow growing massive corals,” Ecol. Eng., vol. 123, pp. 86–94, 2018.

[41] E. Chornesky, “The ties that bind: inter-clonal cooperation may help a fragile coral dominate shallow high-energy reefs,” Mar. Biol., vol. 109, no. 1, pp. 41–51, 1991.

[42] U. Frank and B. Rinkevich, “Nontransitive patterns of historecognition phenomena in the Red Sea hydrocoral *Millepora dichotoma*,” Mar. Biol., vol. 118, no. 4, pp. 723–729, 1994.

[43] Y. Nozawa and Y. Loya, “Genetic relationship and maturity state of the allorecognition system affect contact reactions in juvenile *Seriatopora* corals,” Mar. Ecol. Prog. Ser., vol. 286, pp. 115–123, 2005.

[44] D. Gateno, A. Israel, Y. Barki et al., “Gastrovascular circulation in an octocoral: evidence of significant transport of coral and symbiont cells,” Biol. Bull., vol. 194, no. 2, pp. 178–186, 1998.

[45] E. H. Gladfelter, “Circulation of fluids in the gastrovascular system of the reef coral *Acropora cervicornis*,” Biol. Bull., vol. 165, no. 3, pp. 619–636, 1983.

[46] E. Titlyanov, T. Titlyanova, V. Leletkin et al., “Degradation of zooxanthellae and regulation of their density in hermatypic corals,” Mar. Ecol. Prog. Ser., vol. 139, pp. 167–178, 1996.

[47] J. Stimson and R. A. Kinzie III, “The temporal pattern and rate of release of zooxanthellae from the reef coral *Pocillopora damicornis* (Linnaeus) under nitrogen-enrichment and control conditions,” J. Exp. Mar. Biol. Ecol., vol. 153, no. 1, pp. 63–74, 1991.

[48] M. Hirose, H. Yamamoto, and M. Nonaka, “Metamorphosis and acquisition of symbiotic algae in planula larvae and primary polyps of *Acropora* spp.,” Coral Reefs, vol. 27, no. 2, pp. 247–254, 2008.

[49] H. J. Scharfenstein, W. Y. Chan, P. Buerger et al., “Evidence for de novo acquisition of microalgal symbionts by bleached adult corals,” ISME J., vol. 16, no. 6, pp. 1676–1679, 2022.

[50] J. Schindelin, I. Arganda-Carreras, E. Frise et al., “Fiji: an open-source platform for biological-image analysis,” Nat. Methods, vol. 9, no. 7, pp. 676–682, 2012.

[51] Y. Li, X. Liao, X. Wang et al., “Polyp-canal reconstruction reveals evolution toward complexity in corals,” Research, vol. 6, p. 0166, 2023.

[52] N. Rädecker, C. Pogoreutz, H. M. Gegner et al., “Heat stress destabilizes symbiotic nutrient cycling in corals,” Proc. Natl. Acad. Sci., vol. 118, no. 5, p. e2022653118, 2021.

[53] D. Yellowlees, T. A. V. Rees, and W. Leggat, “Metabolic interactions between algal symbionts and invertebrate hosts,” Plant, Cell Environ., vol. 31, no. 5, pp. 679–694, 2008.

[54] M. Pernice, A. Meibom, A. van den Heuvel et al., “A single-cell view of ammonium assimilation in coral–dinoflagellate symbiosis,” ISME J., vol. 6, no. 7, pp. 1314–1324, 2012.

[55] O. H. Shapiro, V. I. Fernandez, M. Garren et al., “Vortical ciliary flows actively enhance mass transport in reef corals,” Proc. Natl. Acad. Sci., vol. 111, no. 37, pp. 13391–13396, 2014.

[56] C. O. Pacherres, S. Ahmerkamp, G. M. Schmidt-Grieb et al., “Ciliary vortex flows and oxygen dynamics in the coral boundary layer,” Sci. Rep., vol. 10, no. 1, p. 7541, 2020.

[57] T. Bouderlique, J. Petersen, L. Faure et al., “Surface flow for colonial integration in reef-building corals,” Curr. Biol., vol. 32, no. 12, pp. 2596–2609.e7, 2022.

[58] S. A. Selvan, C. O. Pacherres, M. Kühl et al., “Unravelling three-dimensional active transport by ciliary arrays on coral surfaces,” PRX Life, vol. 4, no. 2, p. 023020, 2026.

[59] A. P. Parrin, T. L. Goulet, M. A. Yaeger et al., “*Symbiodinium* migration mitigates bleaching in three octocoral species,” J. Exp. Mar. Biol. Ecol., vol. 474, pp. 73–80, 2016.

[60] J. A. Schwarz, D. A. Krupp, and V. M. Weis, “Late larval development and onset of symbiosis in the scleractinian coral *Fungia scutaria*,” Biol. Bull., vol. 196, no. 1, pp. 70–79, 1999.

[61] T. Bockel and B. Rinkevich, “Rapid recruitment of symbiotic algae into developing scleractinian coral tissues,” J. Mar. Sci. Eng., vol. 7, no. 9, p. 306, 2019.

