## Supplementary Information for "Stronger together: isograft fusion accelerates recovery and may facilitate heat-evolved symbiont transmission in bleached adult corals"

**Materials and Methods**

*S1: DNA extraction, PCR amplification and library preparation*

*DNA extraction*

DNA extractions followed the methodology of Wilson et al. (2002). Briefly, coral tissue samples (collected as described above) (~2 mm³, see below) were lysed and digested using bead-beating (Sigma G1152 - 100G, 710-1180 µm), lysozyme (67.5 µg/mL), and proteinase K (535 µg/mL). The resulting homogenate was incubated at 65°C for 90 minutes. Following incubation, 250 µL of potassium acetate (KOAc) was added per 1 mL of homogenate. Samples were thoroughly mixed, incubated on ice for 30 minutes, and centrifuged at 25,000×g for 15 minutes at 4°C. The supernatant was then carefully decanted, and DNA precipitation was achieved by adding 0.8 volumes of isopropanol. Genomic DNA was collected by centrifugation at 25,000×g for 12 minutes at room temperature, then washed with 200 µL of 70% ethanol. The supernatant was discarded, and the DNA pellet was air-dried for 20–30 minutes. Finally, the DNA pellet was resuspended in ~35 µL of Milli-Q water and stored at 4°C.

*Amplification*
Prior to amplification, 10 µL of eluted DNA from each technical replicate was combined into a single pooled sample. PCR amplification of the Internal Transcribed Spacer 2 (ITS2) region was performed in triplicate using the SYM_VAR_5.8S2 (forward: 5’-GTGACCTATGAACTCAGGAGTCGAATTGCAGAACTCCGTGAACC-3’) and SYM_VAR_REV (reverse: 5’-CTGAGACTTGCACATCGCAGCCGGGTTCWCTTGTYTGACTTCATGC-3’) primers developed by Hume *et al*. 2018 [10] with Illumina adapters underlined. The PCR reactions were set up with 30 µl of Taq PCR Master Mix (201445, Qiagen, Hilden, Germany), 1 µl of 1:10 diluted DNA template, 1.5 µl of forward and reverse primer (10 µM working solution), and 24 µl of Milli-Q ultrapure water. The optimised thermocycling protocol consisted of an initial denaturation step at 95.0°C for 15 min, followed by 18 cycles of denaturation at 95°C for 15 s, annealing at 56°C, and extension at 72°C for 30 s each, with a final extension step at 72°C for 7 min. The triplicate PCR reactions then underwent an indexing PCR reaction using the Nextera XT Library Prep kit (Illumina) with a dual indexing strategy. Indexed triplicate PCR amplicons were then pooled and cleaned using AMPure XP beads following the manufacturer’s protocol. Product quantity and size were visualized on 1% agarose 1 x TAE agarose gel.

*S2:* *SymPortal Analysis*

Demultiplexed raw sequences were submitted to SymPortal for ITS2 profiling. SymPortal identifies ITS2 profiles based on recurring assemblages of intragenomic variants across samples, representing putative Symbiodiniaceae taxa. Replicates with a single defining intragenomic variant (DIV) were excluded due to insufficient sequence depth. DNA sequence reads ≤650 were removed from all samples to eliminate background contamination. Samples with a total read count of less than 5,000 were subsequently excluded. Sequencing data were downloaded from SymPortal and analysed in R under version 4.2.0 (R Core Team, 2022).

**Results**

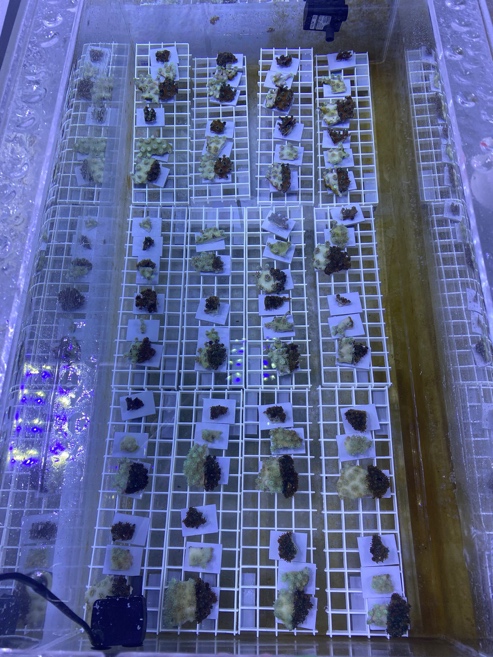

**Fig S1** Experimental setup of *Galaxea fascicularis* in a 100 L tank containing both fused and unfused donor–recipient pairs. Corals assigned to the symbiont exclusion (EC) treatment were housed separately and are not shown in this tank.

**Table S1** Days required for succesful fusion and tissue integration to be observed for each donor-recipient fusion pairing. Fusion was considered to have occured when the tissues of adjoining fragments had visibly merged in at least once location at the boundary and tissue origin was no longer distinguishable.

| **Species** | **Colony** | **Fusion pairing** | **Days required for succesful fusion** |
| --- | --- | --- | --- |
| *Galaxea fascicularis* | gA | 1845_2-1845_9 | 3 |
| *Galaxea fascicularis* | gA | 1845_3-1845_15 | 3 |
| *Galaxea fascicularis* | gA | 1888_2-1888_3 | 3 |
| *Galaxea fascicularis* | gA | 1888_1-1888_4 | 3 |
| *Galaxea fascicularis* | gA | 1888_13-1888_13 | 5 |
| *Galaxea fascicularis* | gA | 1915_1-1915_10 | 5 |
| *Galaxea fascicularis* | gA | 1915_3-1915_13 | 3 |
| *Galaxea fascicularis* | gA | 1915_2-1915_X | 3 |
| *Galaxea fascicularis* | gB | 1623_4-1623_15 | 5 |
| *Galaxea fascicularis* | gB | 1635_2-1635_5 | 6 |
| *Galaxea fascicularis* | gB | 1635_4-1635_13 | 5 |
| *Galaxea fascicularis* | gB | 1642_2-1642_1 | 5 |
| *Galaxea fascicularis* | gB | 1642_1-1642_? | 4 |
| *Galaxea fascicularis* | gB | 1642_4-1642_2 | 5 |
| *Galaxea fascicularis* | gB | 1669_3-1669_9 | 5 |
| *Galaxea fascicularis* | gB | 1669_1-1669_11 | 5 |
| *Galaxea fascicularis* | gC | 1282_2-1282_3 | 3 |
| *Galaxea fascicularis* | gC | 1313_1-1313_10 | 3 |
| *Galaxea fascicularis* | gC | 1321_1_D-1321_1 | 4 |
| *Galaxea fascicularis* | gC | 1321_D-1321_6 | 4 |
| *Galaxea fascicularis* | gC | 1231_3-1321_7 | 3 |
| *Galaxea fascicularis* | gC | 1282_3-1328_10 | 4 |
| *Galaxea fascicularis* | gC | 1328_1-1328_13 | 3 |
| *Galaxea fascicularis* | gC | 1282_1-1328_15 | 3 |
| *Psammocora columna* | pA | 16-1 | 9 |
| *Psammocora columna* | pA | 17-2 | 9 |
| *Psammocora columna* | pA | 20-3 | 9 |
| *Psammocora columna* | pA | 21-5 | 9 |
| *Psammocora columna* | pA | 22-4 | 9 |
| *Psammocora columna* | pA | 23-6 | 9 |
| *Psammocora columna* | pA | 28-9 | 9 |
| *Psammocora columna* | pB | 52-31 | 9 |
| *Psammocora columna* | pB | 45-38 | 11 |
| *Psammocora columna* | pB | 47-39 | 9 |
| *Psammocora columna* | pB | 48-40 | 11 |
| *Psammocora columna* | pB | 50-41 | 9 |
| *Psammocora columna* | pB | 51-43 | 9 |
| *Psammocora columna* | pB | 49-44 | 9 |
| *Psammocora columna* | pC | 70-58 | 9 |
| *Psammocora columna* | pC | 72-59 | 9 |
| *Psammocora columna* | pC | 73-63 | 9 |
| *Psammocora columna* | pC | 75-65 | 9 |
| *Psammocora columna* | pC | 74-66 | 9 |
| *Psammocora columna* | pC | 81-68 | 9 |
| *Psammocora columna* | pC | 76-69 | 9 |
| *Psammocora columna* | pD | 100-85 | 9 |
| *Psammocora columna* | pD | 101-86 | 9 |
| *Psammocora columna* | pD | 102-87 | 10 |
| *Psammocora columna* | pD | 103-88 | 11 |
| *Psammocora columna* | pD | 104-89 | 13 |
| *Psammocora columna* | pD | 105-92 | 13 |
| *Psammocora columna* | pD | 106-99 | 15 |

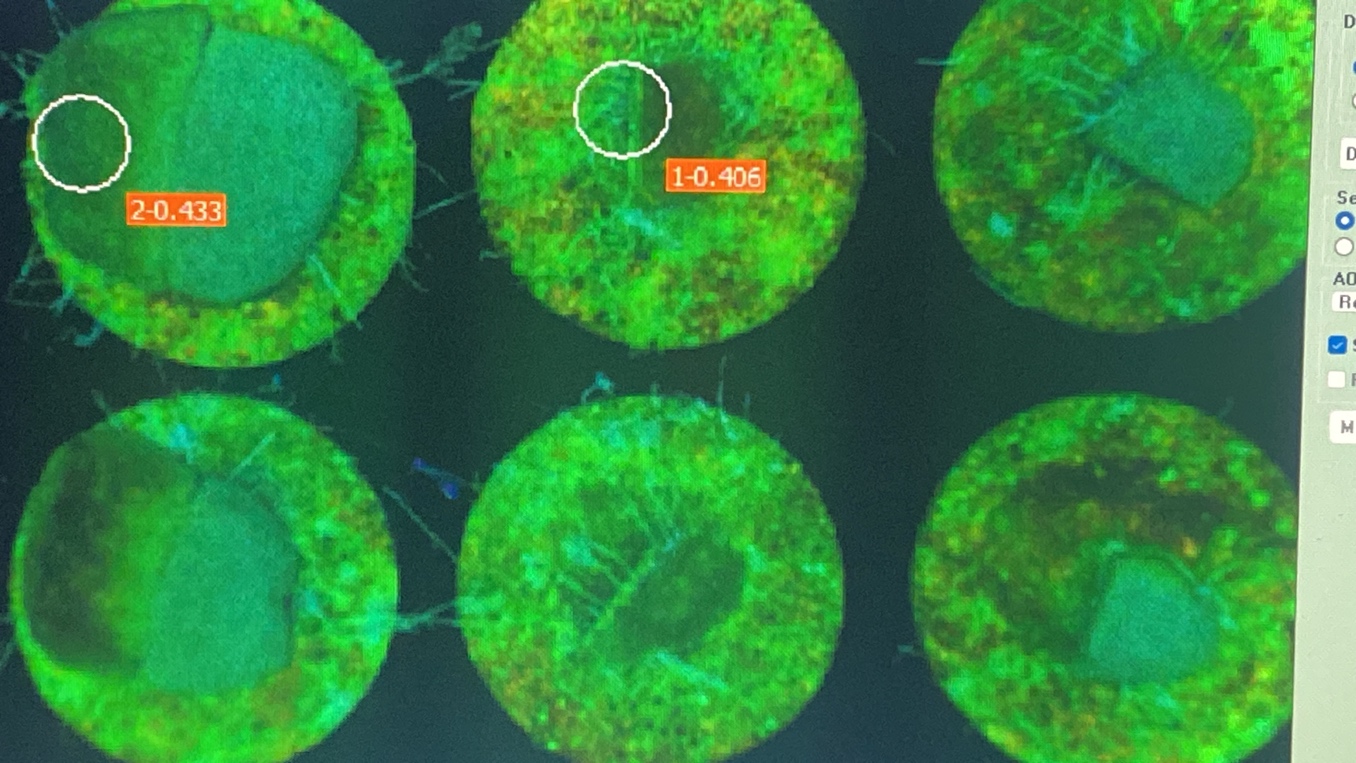

**Fig S2** Representative image of a fused *Psammocora columna* pair, showing the symbiont-rich donor fragment on the left (blue-green) and the bleached recipient fragment on the right. The image depicts spatial variation in photochemical efficiency across the fusion boundary, as visualised using PAM fluorescence imaging. Red scale bar = 1 cm.

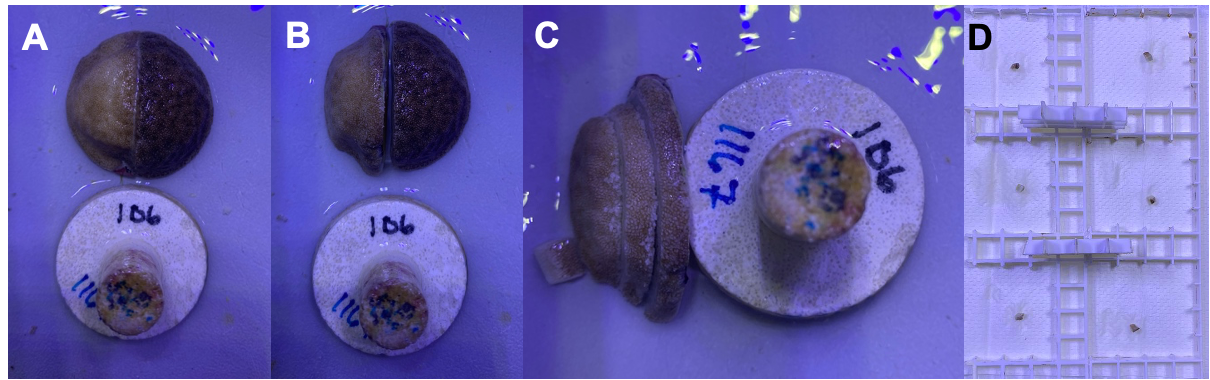
**Fig S3** Workflow for tissue extraction from fused donor–recipient coral pairs. **(A)** Fused pair is first removed from the plug. **(B)** Donor and recipient fragments are then separated along the fusion line. **(C)** The recipient fragment is subdivided into two regions: boundary (adjacent to the fusion line, following a ~1 mm tissue shave) and far (distal from the fusion site). Donor subdivision not shown. **(D)** Generation of triplicate tissue samples from each region: recipient far (left) and recipient boundary (right).

**Table S2** Instances of SS8 DIV occurrences by category combination in *Galaxea fascicularis*

| Colony | Donor*Recipient*HT | Donor*Recipient | Donor only | Recipient only |
| --- | --- | --- | --- | --- |
| gA | 5/6 | 1/6 | 0/6 | 0/6 |
| gB | 5/7 | 0/7 | 1/7 | 1/7 |
| gC | 3/6 | 1/6 | 2/6 | 0/6 |

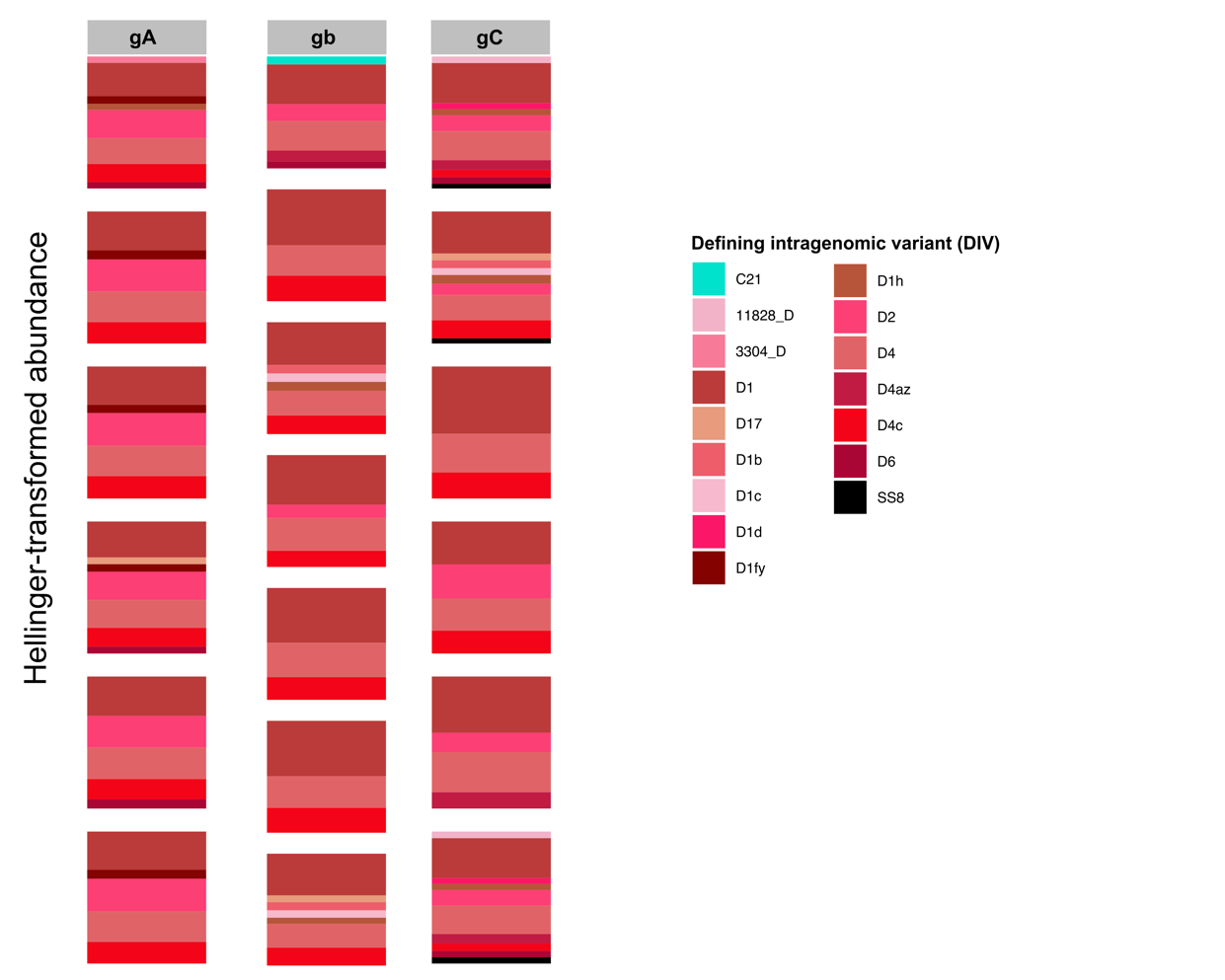

**Fig S4** *Galaxea fascicularis* ITS2 Symbiodiniaceae community composition in experimental donor fragments prior to the start of the experiment (0 DPF). Bar plots show Hellinger-transformed ITS2-derived symbiont communities for three *G. fascicularis* genotypes (gA, gB, and gC). Each panel represents an individual fragment, with bars indicating the proportional abundance of dominant intragenomic variants (DIVs), coloured by genus or identified as SS8 (black).

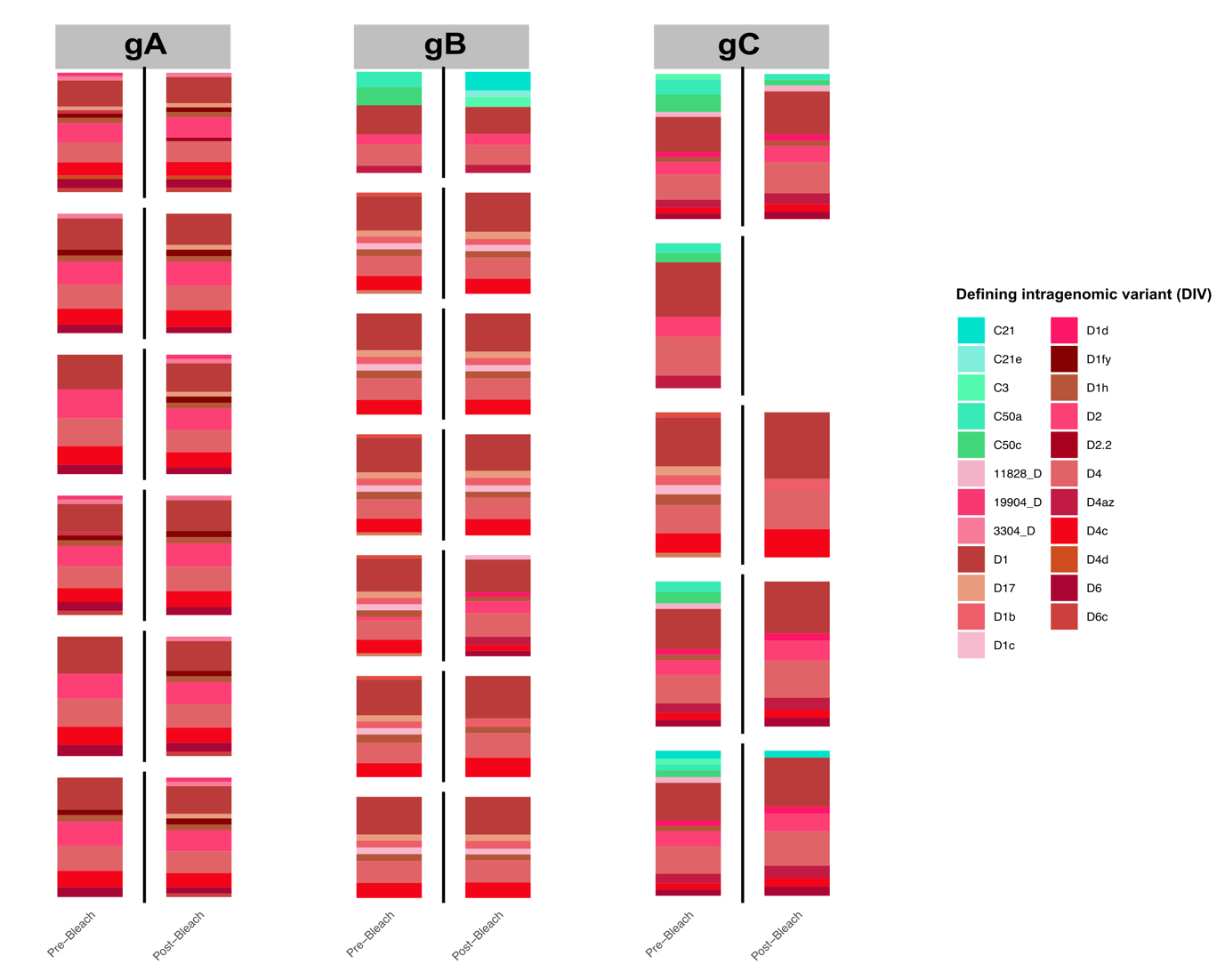
**Fig. S5.** *Galaxea fascicularis* ITS2 Symbiodiniaceae community composition in experimental recipient fragments pre and post chemical bleaching. Bar plots show Hellinger-transformed ITS2-derived symbiont communities for three *G. fascicularis* genotypes (gA, gB, and gC). Each panel represents an individual fragment, with bars indicating the proportional abundance of dominant intragenomic variants (DIVs), coloured by genus.
